# An extended Tudor domain within Vreteno interconnects Gtsf1L and Ago3 for piRNA biogenesis in *Bombyx mori*

**DOI:** 10.1101/2023.03.23.533951

**Authors:** Alfred W. Bronkhorst, Chop Y. Lee, Martin M. Möckel, Sabine Ruegenberg, Antonio M. de Jesus Domingues, Shéraz Sadouki, Tetsutaro Sumiyoshi, Mikiko C. Siomi, Lukas Stelzl, Katja Luck, René F. Ketting

**Author notes:** Shared correspondence.

## Abstract

Piwi-interacting RNAs (piRNAs) direct PIWI proteins to transposons to silence them, thereby preserving genome integrity and fertility. The piRNA population can be expanded in the ping-pong amplification loop. Within this process, piRNA-associated PIWI proteins (piRISC) enter the nuage to cleave target RNA, which is stimulated by Gtsf proteins. The resulting cleavage product gets loaded into an empty PIWI protein to form a new piRISC complex. However, for piRNA amplification to occur, it is required that new RNA substrates, Gtsf-piRISC and empty PIWI proteins are all in physical proximity. In this study we show that BmGtsf1L binds to piRNA-loaded BmAgo3 and co-localizes to BmAgo3-BmVreteno positive granules. Biochemical assays further revealed that conserved residues within the unstructured tail of BmGtsf1L directly interact with BmVreteno. Using a combination of AlphaFold modeling, atomistic molecular dynamics simulations and in vitro assays we identified a novel binding interface on a BmVreteno-eTudor domain, which is required for BmGtsf1L binding. Our study reveals that a single eTudor domain within BmVreteno provides two binding interfaces and thereby interconnects piRNA-loaded BmAgo3 and BmGtsf1L.

## Introduction

Animal germ cells utilize the Piwi-interacting (pi)RNA pathway as a mechanism to silence transposons, thereby maintaining genome stability and fertility (Czech *et al*, 2018; Ozata *et al*, 2019). A defective piRNA pathway leads to transposon de-repression, DNA damage, gametogenesis defects and sterility. The piRNA-pathway can also have non-transposon targets, such as in the silk moth *Bombyx mori* where piRNAs regulate sex determination (Kiuchi *et al*, 2014). piRNAs are about 24-31 nucleotides in size and associate with PIWI-clade Argonaute proteins to guide them to complementary targets (Ghildiyal & Zamore, 2009). Therefore, piRNAs are a key, specificity-determining component of the Piwi pathway.

In *Drosophila* germ cells, precursor piRNAs (pre-pre-piRNAs) are transcribed within large dual-strand clusters and are exported from the nucleus for subsequent processing within the cytoplasm (Brennecke *et al*, 2007; Ozata *et al*, 2019; Czech *et al*, 2018). piRNAs can be loaded into different PIWI proteins. *Drosophila* expresses three PIWI proteins, Piwi, Aubergine (Aub) and Ago3. Piwi and Aub are predominantly loaded with antisense piRNAs, whereas Ago3 mostly incorporates sense piRNAs (Brennecke *et al*, 2007; Gunawardane *et al*, 2007). In the silkworm only two cytoplasmic PIWI proteins are expressed, Siwi and BmAgo3 (Kawaoka *et al*, 2009). Siwi-associated piRNAs are mostly antisense and are responsible for the cleavage of sense transposon mRNA, whereas BmAgo3 binds sense piRNAs and triggers antisense piRNA precursor cleavage.

The current model for piRNA biogenesis suggests that cytoplasmic piRNA processing occurs by two interconnected mechanisms. One step of piRNA processing takes place within the nuage, a germline-specific phase-separated structure that surrounds the nuclear membrane. Here, piRNA-guided (called trigger piRNA) endonuclease activity of one PIWI protein (e.g. Ago3) generates the 5’ monophosphate end of a complementary piRNA precursor transcript (Homolka *et al*, 2015; Han *et al*, 2015; Mohn *et al*, 2015; Gainetdinov *et al*, 2018). This so-called responder pre-piRNA is subsequently incorporated into an unloaded PIWI protein (*Drosophila* Aub or silkworm Siwi), which is still too long at its 3’-end. For further pre-piRNA processing the PIWI protein then migrates to the mitochondrial outer membrane. Here, the second step of piRNA processing is mediated by the endonuclease Zucchini (Zuc), which mediates responder pre-piRNA 3’-end formation by cleaving 5’ to an available uridine (Mohn *et al*, 2015; Han *et al*, 2015). Pre-piRNA 3’-end resection is further completed by trimming and methylation to generate a mature piRISC complex (Izumi *et al*, 2016; Kawaoka *et al*, 2011; Hayashi *et al*, 2016; Horwich *et al*, 2007). The mature piRISC complex is liberated from the mitochondria into the cytosol to cleave complementary target RNA, resulting in its degradation. Alternatively, the mature piRISC complex (Aub or Siwi) can transit back to the nuage (now serving as a trigger piRNA) to bind complementary target RNA and to initiate a new round of PIWI-catalyzed responder piRNA biogenesis leading to new piRISC formation. This consecutive and continuous process of responder and trigger piRNA production, which requires reciprocal cleavages by two paired PIWI proteins, is called the piRNA amplification cycle (or ping-pong loop). Thus, mature piRNAs within the ping-pong cycle are generated by the combined action of PIWI-slicing and Zuc-cleavage.

Notably, Zuc-mediated processing of the responder pre-piRNA 3’-end simultaneously generates the 5’-end of a new pre-piRNA substrate for phased piRNA biogenesis, which results in the production of trailer piRNAs (Mohn *et al*, 2015; Han *et al*, 2015). Initiator and responder piRNAs that are generated via the ping-pong cycle increase the abundance of an existing pool of piRNAs, whereas the Zuc-dependent trailer piRNAs expand the repertoire of piRNA sequences. In *Drosophila*, phased piRNAs predominantly associate with Piwi and translocate to the nucleus to induce transcriptional gene silencing through the deposition of repressive chromatin marks (Czech *et al*, 2018). Even though trailer piRNAs are produced, the silkworm does not possess a nuclear piRNA-based silencing pathway (Gainetdinov *et al*, 2018; Izumi *et al*, 2020).

Efficient piRNA amplification within the ping-pong cycle requires that PIWI-mediated target cleavage is confined to molecular surroundings that are compatible with an empty PIWI protein receiving one of the cleavage products. It has been suggested that Tudor-domain containing proteins that reside in the nuage can provide such an environment by acting as a molecular scaffold (Chen *et al*, 2011; Siomi *et al*, 2011). Tudor domains that harbor an aromatic cage can bind to symmetrically dimethylated arginine residues (sDMAs) on client proteins but can also establish sDMA-independent protein interactions (Siomi *et al*, 2010; Chen *et al*, 2011). For example, the *Drosophila* Krimper protein makes sure that cleavage products resulting from Aub-slicing are efficiently loaded into empty Ago3 (Webster *et al*, 2015; Sato *et al*, 2015). Krimper binds sDMA-methylated piRISC-Aub via its aromatic cage-containing Tudor domain, whereas an upstream Tudor domain within Krimper establishes the sDMA-independent interaction with empty Ago3 (Sato *et al*, 2015; Webster *et al*, 2015; Huang *et al*, 2021). Likewise, the multi Tudor domain-containing protein Qin also promotes heterotypic ping-pong between piRISC-Aub and empty, unmethylated Ago3 (Zhang *et al*, 2014, 2011). In silkworms, the handover of piRISC-Siwi cleaved target RNA to empty BmAgo3 is mediated by the RNA-helicase Vasa (Xiol *et al*, 2014; Nishida *et al*, 2015). Notably, Vasa contains intrinsically disordered regions that are involved in the formation of phase-separated structures and seems to be the scaffold for nuage formation (Nott *et al*, 2015). Moreover, the Vasa N-terminus is strongly methylated, indicating that multivalent interactions with Tudor domain-containing proteins also contribute to nuage assembly (Kirino *et al*, 2010). Additional studies in silkworm revealed that BmVreteno brings piRNA-loaded BmAgo3 and empty Siwi together via their Tudor domains to allow new piRISC-Siwi formation (Nishida *et al*, 2020). This may involve the dimerization of two BmVreteno isoforms (BmVreteno-Long and -Short), where the BmVreteno-Long isoform anchors the RNA target through its unique RRM domain. Thus, BmVreteno also acts in the ping-pong cycle but has a reverse role compared to Krimper and Qin, as it enforces Siwi loading instead of BmAgo3 loading.

Recently, Arif et al. reported that gametocyte-specific factor proteins (Gtsf) stimulate the catalytic activity of PIWI proteins (Arif *et al*, 2022). Gtsf proteins act in the piRNA pathway in different species, including flies, silkworms and mice (Ipsaro & Joshua-Tor, 2022). In *Drosophila*, Gtsf1 is required for piRNA-mediated transcriptional gene silencing and is not essential for piRNA biogenesis (Dönertas *et al*, 2013; Ohtani *et al*, 2013). In contrast, mouse Gtsf1 is involved in piRNA amplification and associates with the mouse PIWI proteins Miwi2 and Mili (Yoshimura *et al*, 2018). Moreover, mouse Gtsf1 can enhance the piRNA-directed target cleavage of both Mili and Miwi *in vitro*. Likewise, silkworm Gtsf1 associates with Siwi and enhanced slicing activity was found for Siwi but not for Ago3 (Chen *et al*, 2020; Arif *et al*, 2022). We note that the conditional cleavage by PIWI proteins, which is dependent on Gtsf, provides an interesting possibility to restrict target cleavage to conditions in which an empty PIWI protein may be available to accept a cleavage product, and to prevent RNA cleavage in absence of such empty PIWI proteins. However, it is not known how the Gtsf-piRISC complex is brought in physical proximity with empty PIWI and target RNA.

In this study, we show that silkworm Gtsf1-like (BmGtsf1L), a Gtsf1 paralog, binds piRNA-loaded BmAgo3 and this interaction is stimulated by BmVreteno, a protein known to aid Siwi loading following BmAgo3-mediated target cleavage. Surprisingly, we find that BmGtsf1L and BmAgo3 bind to the same eTudor domain of BmVreteno, and using AlphaFold predictions we uncover a novel binding interface on this eTudor domain that additionally accommodates BmGtsf1L binding. Thus, a single eTudor domain within BmVreteno can serve as a molecular scaffold and interconnects BmGtsf1L and piRISC-BmAgo3 to allow efficient target cleavage only within a context that enables Siwi loading.

## Results

### BmGtsf1L associates with piRNA-loaded BmAgo3

A previous study showed that Gtsf1 is involved in piRNA-regulated sex determination and transposon silencing in the silkworm (Chen *et al*, 2020). The role of its paralog, BmGtsf1L, however, remained elusive. Alignment of Gtsf proteins from different species shows that BmGtsf1L possesses the two conserved CHHC-type zinc (Zn) fingers followed by a short C-terminal tail (Fig. S1A). To find potential binding partners of BmGtsf1L, we transiently expressed HA-tagged BmGtsf1L in BmN4 cells and performed an immunoprecipitation followed by quantitative mass-spectrometry (IP/qMS). Interestingly, many of the enriched proteins are known to play a role in piRNA biogenesis, such as BmAgo3, BmVreteno and Siwi (Fig. 1A, Table S1). Next, we transiently expressed HA-BmGtsf1L together with FLAG-tagged BmAgo3, BmVreteno or Siwi and confirmed that these candidates interact with BmGtsf1L, both in the presence or absence of RNA (Fig. 1B). Also endogenous BmAgo3 associated with transiently expressed BmGtsf1L (Fig. 1C).

**Fig 1.**
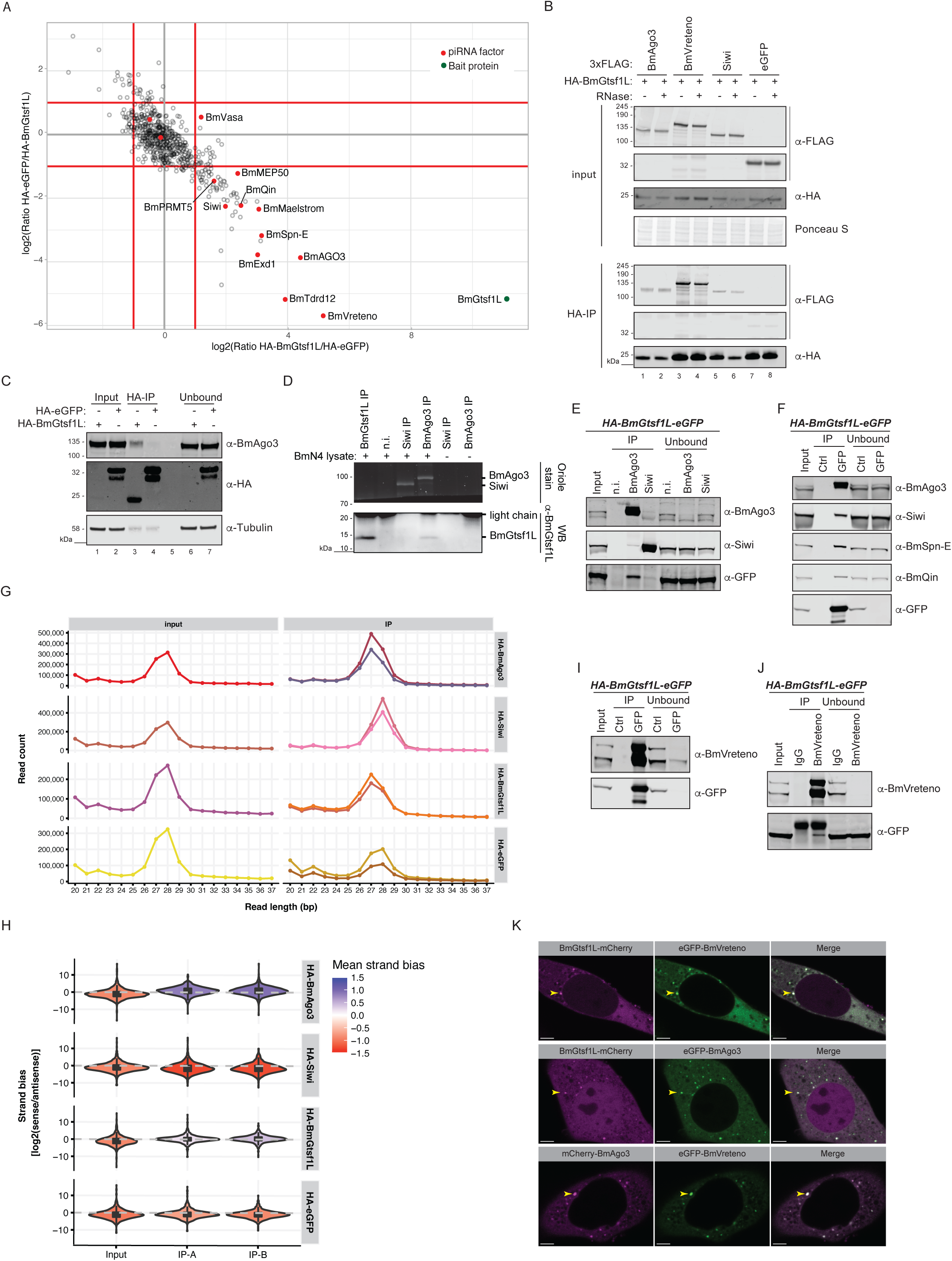
BmGtsf1L associates with piRNA-loaded BmAgo3. a) Anti-HA immunoprecipitation on BmN4 cell lysates where either HA-BmGtsf1L or HA-eGFP was ectopically expressed. The experiment was performed using technical duplicates to perform quantitative mass-spectrometry based detection of peptides using stable dimethyl isotope labeling. Scatterplot showing the log2 converted normalized ratio data for the individual label pairs. The threshold was set to 2-fold enrichment, known piRNA factors are indicated (red dots) as well as the bait protein (BmGtsf1L, green dot). b) Anti-HA immunoprecipitation on BmN4 lysates made from cells that were transfected with the indicated constructs either in presence or absence of RNase A/T1. BmGtsf1L was immunoprecipitated followed by Western blot detection using the indicated antibodies. Expression of 3xFLAG-eGFP served as a negative control. c) Anti-HA immunoprecipitation of HA-BmGtsf1L or HA-eGFP from BmN4 cell lysates followed by immunodetection of endogenous BmAgo3. Anti-tubulin probing served as a loading control. d) Immunoprecipitation using the indicated endogenous antibodies or using non-immune serum (n.i.) as a control in the presence or absence of naïve BmN4 cell lysates. Oriole stain was used to detect the immunopurified BmAgo3 and Siwi complexes, whereas retrieval of endogenous BmGtsf1L was verified by Western blot. e) Immunoprecipitation of endogenous BmAgo3 and Siwi complexes on BmN4 cell lysates from the HA-BmGtsf1L-eGFP stable cell line, followed by Western blot detection using the indicated antibodies. f) GFP (BmGtsf1L) or control (Ctrl) immunoprecipitation on BmN4 cell extracts stably expressing HA-BmGtsf1L-eGFP followed by Western blot detection using the indicated antibodies. g) Small RNA size profiles of input samples and from anti-HA immunopurified complexes. Immunoprecipitations were performed in duplicate on BmN4 cell lysates from cells that were transiently transfected, denoted in the two lines on the right-hand panels. h) Violin plot showing the log2 transformed strand bias of sense to antisense small RNAs from input and IP samples. The mean strand bias is indicated with the color code, where negative and positive values represents antisense and sense bias, respectively. i) GFP (BmGtsf1L) and control immunoprecipitation followed by Western blot detection of endogenous BmVreteno, using an antibody that detects the two BmVreteno isoforms. j) Reciprocal immunoprecipitation using endogenous anti-BmVreteno antibody or rabbit IgG as an isotype control, followed by immunodetection using the indicated antibodies. k) Single-plane confocal micrographs of BmN4 cells co-transfected with BmGtsf1L-mCherry and eGFP-BmVreteno (upper panel), BmGtsf1L-mCherry with eGFP-BmAgo3 (middle panel) or mCherry-BmAgo3 and eGFP-BmVreteno (bottom panel). Yellow triangles indicate a formed granule. Scale bars – 4 µm.

Next, we generated an anti-BmGtsf1L monoclonal antibody, which detected both endogenous as well as epitope-tagged BmGtsf1L (Fig. S1B). Despite the low expression levels of BmGtsf1L, we could detect endogenous BmGtsf1L specifically in BmAgo3 precipitates (Fig. 1D). Unfortunately, the BmGtsf1L antibody was not suitable for immunoprecipitation assays and also did not function in immunostainings. To be able to study BmGtsf1L function in further detail, we generated a BmN4 cell line stably expressing HA-BmGtsf1L-eGFP. Using this stable cell line, we confirmed that BmGtsf1L is mostly enriched in BmAgo3 IPs and hardly in Siwi IPs (Fig. 1E). *Vice versa*, BmAgo3 and Siwi were both co-precipitated with BmGtsf1L and again we observed a stronger enrichment for BmAgo3 (Fig. 1F). Using stable cell lines expressing epitope-tagged PIWI proteins (Fig. S1C), we could show that increased affinity of BmAgo3 for BmGtsf1L was not due to differences in PIWI antibody specificities (Fig. S1D). Moreover, BmSpn-E and BmQin were also co-purified by BmGtsf1L, confirming our initial IP/qMS hits (Fig. 1A, 1F).

To reveal the loading status of endogenous BmAgo3 that associates with BmGtsf1L, we performed BmGtsf1L immunoprecipitation followed by small RNA sequencing. BmGtsf1L-associated small RNA profiles resembled those of BmAgo3 bound small RNAs; showing a clear, defined peak of 27 nucleotides in size, a strong sense-bias, and enrichment for adenine at the 10^th^ position (Fig. 1G, 1H & S1E). Together, these results indicate that BmGtsf1L associates with piRNA-loaded BmAgo3.

Interestingly, BmVreteno has been shown to also interact with piRNA-loaded BmAgo3, and to do so as a dimer (Nishida *et al*, 2020). Using an independently generated anti-BmVreteno antibody that also detects the BmVreteno-Long (L) and BmVreteno-Short (S) isoforms (Fig. S1F, S1G, S1H), we confirm that BmVreteno retrieves BmAgo3 (Fig. S1I). Consistently, both BmVreteno isoforms were also found in BmAgo3 precipitates (Fig. S1J). Next, we assessed the interaction between BmGtsf1L and endogenous BmVreteno. This revealed that BmGtsf1L also brings down both isoforms of endogenous BmVreteno (Fig. 1I). Likewise, BmGtsf1L is also co-precipitated by BmVreteno (Fig. 1J). Together, these data suggest that BmVreteno, BmGtsf1L and piRNA-loaded BmAgo3 form a complex.

### BmGtsf1L resides in BmAgo3 bodies

Nishida et al. recently described that the formation of BmAgo3 bodies is dependent on BmVreteno (Nishida *et al*, 2020). In line with this, we observed co-localization of BmAgo3 and BmGtsf1L with BmVreteno in granules (Fig. 1K, top and lower panels). We note that BmGtsf1L was present in BmAgo3-marked granules, even though the majority of BmGtsf1L was distributed throughout the nucleus and cytoplasm (Fig. 1K, middle panel). Interestingly, additional expression of epitope-tagged BmVreteno, but not of BmAgo3, leaves no detectable expression of BmGtsf1L within the nucleus (Fig. 1K, top and middle panels), indicating that BmVreteno is the major determinant of BmGtsf1L localization, and not BmAgo3.

### Interdependence of BmVreteno-BmAgo3-BmGtsf1L interaction

Next, we assessed the consequences of BmGtsf1L knockdown on the interactions we described above. Knockdown of BmGtsf1L followed by BmVreteno immunoprecipitation revealed that the interaction between BmVreteno and BmAgo3 does not require BmGtsf1L (Fig. 2A). *Vice versa*, both isoforms of BmVreteno were also still retrieved by BmAgo3 following BmGtsf1L depletion (Fig. 2B). These results indicate that the interaction between BmVreteno and BmAgo3 does not require BmGtsf1L.

**Fig 2.**
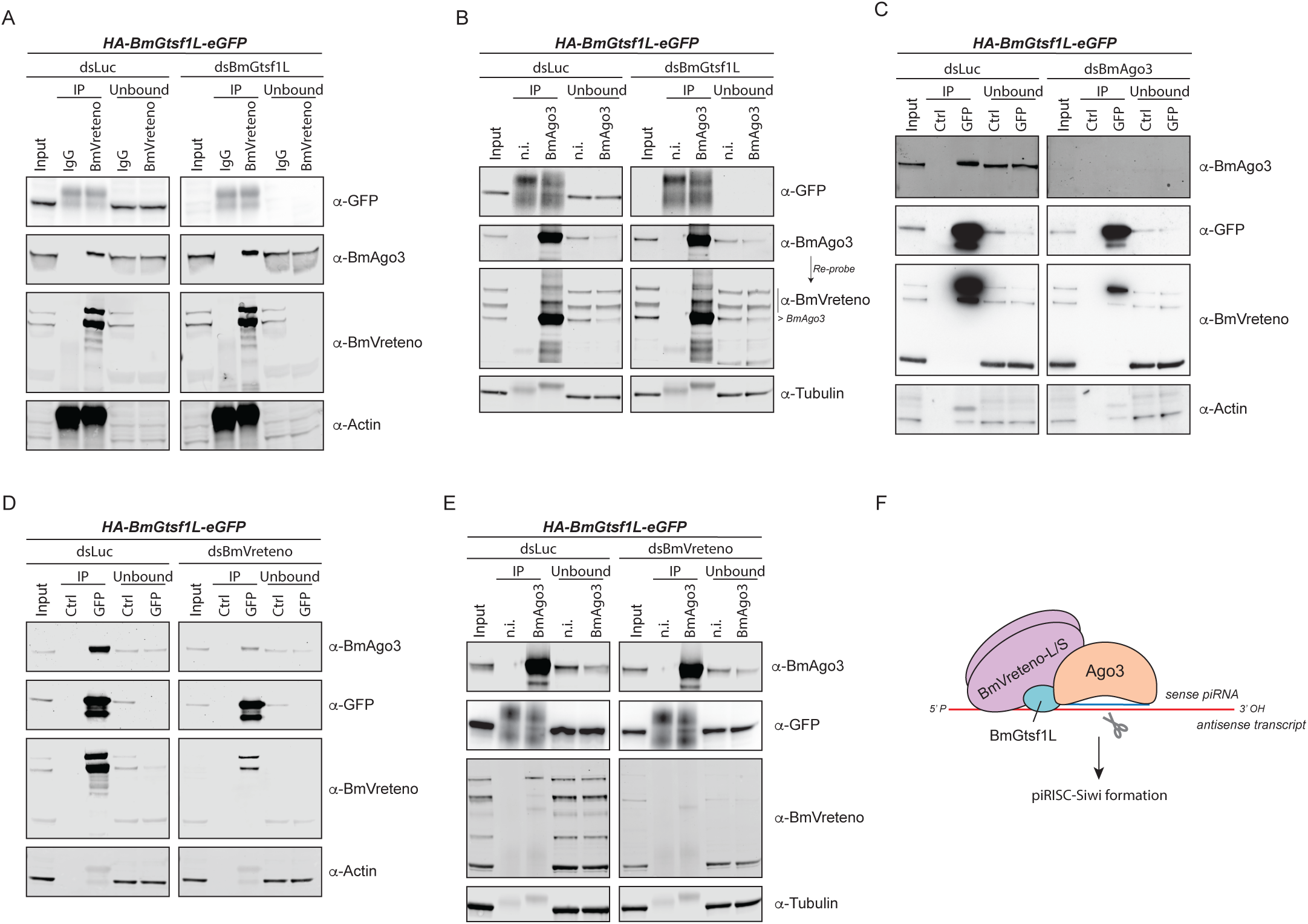
Interdependence of BmVreteno-BmAgo3-BmGtsf1L interaction. dsRNA-mediated gene depletion on BmN4 cells, which stably express HA-BmGtsf1L-eGFP, followed by immunoprecipitation and Western blot detection. a) Luciferase (dsLuc) control knockdown and BmGtsf1L (dsBmGtsf1L) depletion followed by IgG control or anti-BmVreteno immunoprecipitations and detection of retrieved proteins by Western blot using the indicated antibodies. Anti-actin detection served as the loading control. b) Knockdown in BmN4 cells as described in panel (a), followed by non-immune (n.i.) serum control IP or anti-BmAgo3 IP and detection of retrieved proteins by Western blot. Anti-tubulin probing was used as a loading control. c) Knockdown of luciferase (dsLuc) or BmAgo3 (dsBmAgo3) in BmN4 cells. Immunoprecipitation using GFP (BmGtsf1L) or control (Ctrl) magnetic beads, followed by Western blot detection using the indicated antibodies. Anti-actin probing was performed as a loading control. d) Immunoprecipitation and Western blot detection was performed as described in panel (c) but on BmN4 cell extracts from which endogenous BmVreteno was depleted by dsRNA transfection. e) Knockdown of endogenous BmVreteno followed by immunoprecipitation using anti-BmAgo3 antibodies or non-immune serum as a control. Western blot detection of precipitated proteins was performed using the indicated antibodies and detection of anti-tubulin served as a loading control. f) Model on the interconnection between BmGtsf1L, BmAgo3 and BmVreteno. The majority of BmGtsf1L is found in complex with BmVreteno. BmVreteno can stimulate the BmGtsf1L-BmAgo3 interaction, whereas BmAgo3 also fosters the BmGtsf1L-BmVreteno interaction. BmVreteno exist as a Long (L) and Short (S) isoform, which can interact with each other (Nishida *et al*, 2020) and is therefore schematically depicted as a heterodimer.

To test whether BmAgo3 affects the interaction between BmGtsf1L and BmVreteno, we analyzed their association following BmAgo3 knockdown, which was very efficient. This revealed that the interaction between BmGtsf1L and BmVreteno was reduced, but not fully abrogated (Fig. 2C). Given that BmAgo3 was not detected in the IP, these results suggest that BmAgo3 enhances, but is not essential for the interaction between BmGtsf1L and BmVreteno. Finally, we tested the effects of BmVreteno depletion on the BmAgo3-Gtsf1L interaction. While the dsRNA treatment strongly affected BmVreteno levels, we were not able to fully eliminate it, as evidenced by its presence in the BmGtsf1L IPs. Nevertheless, RNAi against BmVreteno strongly diminished the amount of BmAgo3 that was brought down by BmGtsf1L (Fig. 2D), indicating that BmVreteno stimulates the interaction between BmGtsf1L and BmAgo3. We note, however, that in reciprocal BmAgo3 immunoprecipitations small amounts of BmGtsf1L could be retrieved, independent of BmVreteno knock-down status (Fig. 2E). In this experiment no residual BmVreteno was detected in the IPs. We conclude that a small fraction of BmGtsf1L binds BmAgo3 independent of BmVreteno, but that the majority of BmGtsf1L is found in complex with BmVreteno and that this stimulates the BmGtsf1L-BmAgo3 interaction (Fig. 2F).

### BmAgo3 interacts with BmVreteno eTD1 via methylated arginine residues

BmVreteno contains two Tudor domains (TDs) that are confidently predicted by Pfam and SMART Hidden Markov Models (HMMs) (Letunic *et al*, 2021; Mistry *et al*, 2021). A third match of the Pfam Tudor HMM around residue 500 exceeded the e-value threshold, and thus was not significant. Alignment of the two confidently predicted Tudor domains to those of *Drosophila* Tudor-SN and Tudor-eTD11, for which crystal structures have been resolved (Liu *et al*, 2010; Friberg *et al*, 2009), indicates that BmVreteno TD1 contains an aromatic cage (Fig. S2A). It is well known that aromatic cages within TD domains can bind to methylated arginines of client proteins and thereby mediate protein-protein interactions (Siomi *et al*, 2010; Chen *et al*, 2011). When co-expressing BmVreteno with a BmAgo3 variant that cannot be methylated at its N-terminus (5RK), we lost interaction between both (Fig S2B), suggesting that arginine-methylation is a prerequisite for its association with BmVreteno. This is in line with previous work, which showed that the aromatic cage of TD1 is involved in BmAgo3 interaction (Nishida *et al*, 2020).

In addition, a BmAgo3 piRNA-loading defective mutant (YK>LE) also revealed a strong loss of interaction with BmVreteno (Fig. S2B), which is in line with the observation that unloaded BmAgo3 does not co-localize with BmVreteno and fails to form BmAgo3 bodies (Nishida *et al*, 2020). This could indicate that BmAgo3 becomes methylated only following piRNA-binding,which has been observed for *Drosophila* Aubergine (Huang *et al*, 2021; Webster *et al*, 2015). Notably, a BmAgo3 slicing mutant (DADH) does not show loss of interaction with endogenous BmVreteno (Fig. S2B). Taken together, our data, combined with the findings of Nishida et al. (Nishida *et al*, 2020), suggests that the aromatic cage of the BmVreteno TD1 domain mediates the interaction with the methylated N-terminus of BmAgo3.

### The BmGtsf1L C-terminus establishes an interaction with BmVreteno

To understand how BmGtsf1L binds to BmVreteno we tested which region of BmGtsf1L interacted with endogenous BmVreteno. At the same time, we also probed for BmAgo3. A BmGtsf1L fragment missing the N-terminal part, including the two CHHC Zn fingers, could still retrieve BmVreteno, as well as BmAgo3 (Fig. 3A, 3B). By contrast, deletion of the likely disordered BmGtsf1L C-terminus completely abolished the interaction with BmVreteno, while it allowed some interaction with BmAgo3 (Fig. 3A, 3B). Studies in *Drosophila* showed that aromatic residues within the C-terminus of BmGtsf1 regulate Piwi binding (Dönertas *et al*, 2013; Ohtani *et al*, 2013). Therefore, we checked for the presence of aromatic residues within BmGtsf1L and studied how mutagenesis of these residues would affect its interaction with either BmAgo3 or BmVreteno. The BmGtsf1L tyrosine residue mutant (Y88A) retrieved BmAgo3 and BmVreteno to a similar extend as wildtype BmGtsf1L, whereas mutation of the conserved tryptophan residue (W99A) affected the BmAgo3 interaction and completely abrogated the interaction with BmVreteno (Fig. 3A, 3C, & Fig. S1A). The BmGtsf1L double point mutant (YW>AA) displayed similar effects when compared to the W99A single mutant. We could also show that BmGtsf1L (W99A) remained uniformly distributed in the nucleus and cytoplasm, even though BmAgo3 granules were still present (Fig. 3D, 3E). We thus identified the conserved tryptophan residue (W99) within the C-terminal tail of BmGtsf1L to be essential for interaction with BmVreteno and to enhance binding to BmAgo3.

**Fig 3.**
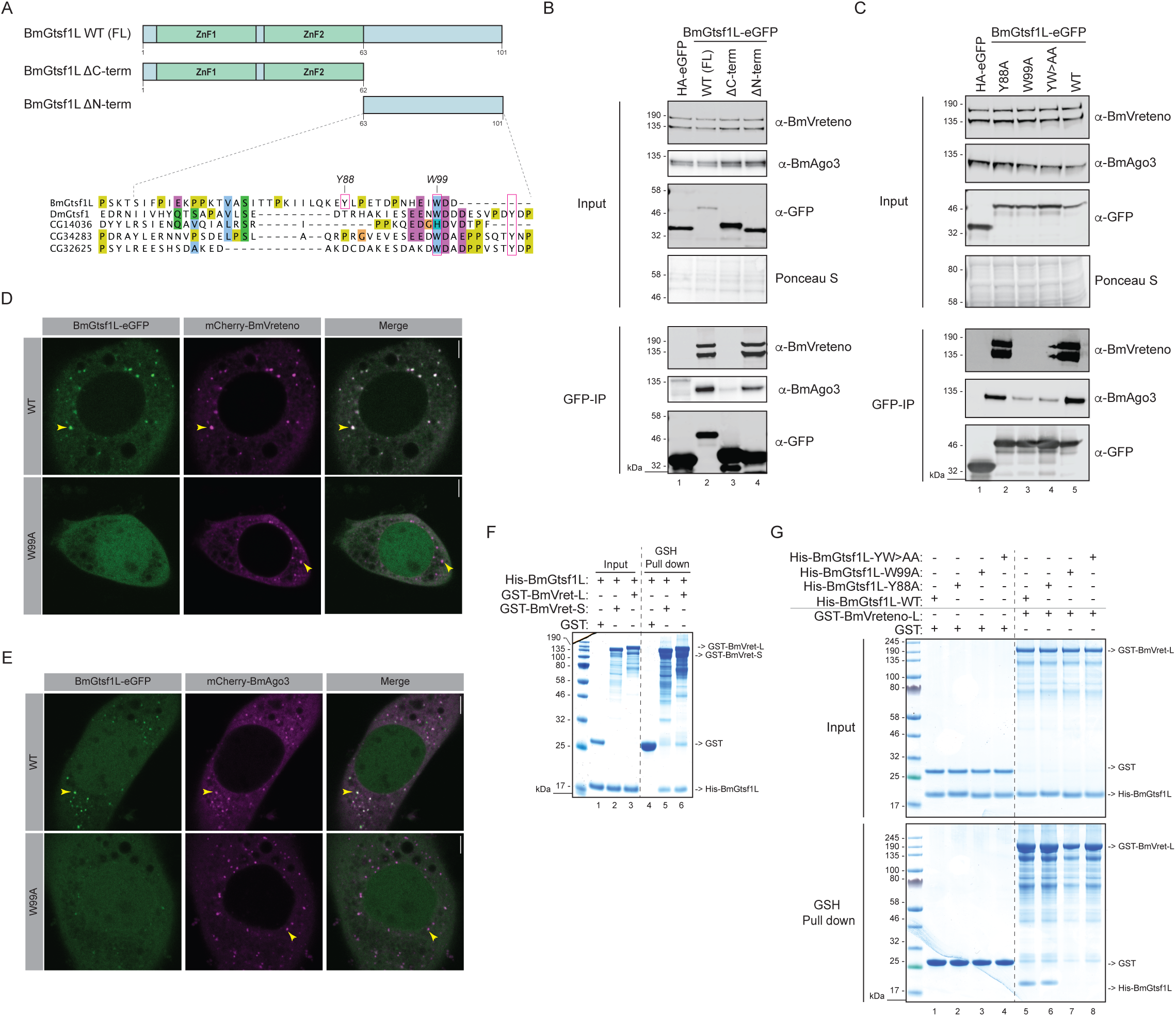
The BmGtsf1L C-terminus establishes a direct interaction with BmVreteno. a) Overview of BmGtsf1L domain architecture with two N-terminally located zinc fingers (ZnF1 and ZnF2, respectively). The two deletion variants used in panel (b) to address binding to BmAgo3 and BmVreteno are also depicted (top). Alignment of BmGtsf1L to GTSF proteins from *Drosophila*. Conserved tryptophan (W) and tyrosine (T) residues of DmGtsf1 that were shown to be involved in Ago3 interaction are boxed in magenta (Dönertas *et al*, 2013). BmGtsf1L contains another aromatic residue (Y88) in its C-terminus (small magenta box), which is not conserved. Clustal Omega alignment was processed with Jalview software (bottom). b) GFP-IP (BmGtsf1L) on BmN4 cell extracts from cells that were transfected with full length (FL) BmGtsf1L-eGFP and with their deletion variants. Transfection of HA-eGFP served as a control. Western blot was performed with indicated antibodies and Ponceau S staining served as a loading control. c) Same as in panel (b) but now BmGtsf1L-eGFP mutants, in which aromatic residues were substituted with alanine, were transfected. d) Single-plane confocal micrographs of BmN4 cells transfected with BmGtsf1L-eGFP wildtype (WT) or the W99A mutant together with mCherry-BmVreteno. Yellow triangles indicate a formed granule. Scale bars – 4 µm. e) Single-plane confocal micrographs of BmN4 cells transfected with BmGtsf1L-eGFP wildtype (WT) or the W99A mutant together with mCherry-BmAgo3. Yellow triangles indicate a formed granule. Scale bars – 4 µm. f) Analysis of the interaction between BmVreteno and BmGtsf1L by GSH pull-down assays. GST alone or GST-BmVreteno L/S was incubated with His-BmGtsf1L. Input and elution fractions were analyzed by SDS-PAGE followed by Coomassie staining. g) I*n vitro* GSH pull-down assay for GST alone or for GST-BmVreteno-L incubated with His-tagged BmGtsf1L variants. Proteins from the SDS-PAGE gel are detected by Coomassie staining.

### BmVreteno directly interacts with BmGtsf1L

The above results prompted us to test the hypothesis that BmVreteno and BmGtsf1L interact directly. Using an *E. coli* expression system we succeeded in the expression and purification of full length BmVreteno-L, BmVreteno-S and BmGtsf1L. Notably, GST-BmVreteno eluted as multimeric proteins from gel filtration columns and associated with nucleic acids (Fig. S3A-S3C). Using these proteins in GSH pull-down assays we could show that BmGtsf1L interacts directly with both isoforms of BmVreteno (Fig. 3F). Furthermore, we could recapitulate the effects of the mutations described above on the BmVreteno-BmGtsf1L interaction *in vitro*: recombinant BmGtsf1L-W99A failed to interact with BmVreteno (Fig. 3G). We conclude that the C-terminal end of BmGtsf1L is sufficient to bind directly to BmVreteno, and that BmGtsf1L-W99 plays a crucial role in this interaction.

### BmGtsf1L binds to BmVreteno eTD1

We next analyzed which region of BmVreteno is involved in its interaction with BmGtsf1L. Using truncation analysis, we found that the C-terminal region of BmVreteno, containing the two PFAM/SMART predicted TDs, was required (Fig. S3D, S3E). However, purification of fragments containing individual predicted TD domains to probe binding with BmGtsf1L failed. To improve fragment design we turned to AlphaFold as a novel artificial intelligence-based tool for protein structure prediction. To our surprise, AlphaFold confidently predicted three extended Tudor domain folds within full length BmVreteno, which are also referred to as TSN folds (Fig. 4A, Suppl Data 1) (Liu *et al*, 2010). Hereafter, we refer to these three domains as AF-eTD0, 1, and 2 where AF-eTD0 corresponds to the newly identified eTD domain. The prediction of this additional Tudor domain is in line with IUPred predictions suggesting that this region is structured (Fig. 4A). It also overlaps with the non-significant Tudor HMM match from SMART/Pfam mentioned earlier. In addition, AlphaFold predicted with high confidence the structures and boundaries of the RRM and MYND domain further upstream of the three eTD domains (Fig. 4B). As indicated by the predicted aligned error matrix, AlphaFold is very uncertain about the relative orientation of the RRM, MYND, and AF-eTD0 domain to the rest of the protein (Fig. 4B). This suggests that they are unlikely to establish intramolecular contacts with other regions in BmVreteno while AlphaFold is somewhat more certain about the predicted relative orientation of AF-eTD1 and AF-eTD2 to each other. Based on the predicted AF-eTD boundaries we designed novel fragments carrying individual eTD domains to test if any of these AF-eTDs mediate the binding to BmGtsf1L. BmN4 cells were transfected with individual HA-eGFP-tagged AF-eTDs of BmVreteno together with mCherry-3xFLAG-BmGtsf1L. Only BmVreteno AF-eTD1 was retrieved in BmGtsf1L immunoprecipitations (Fig. 4C). Likewise, recombinant BmGtsf1L was only co-precipitated with now purifiable GST-tagged BmVreteno-AF-eTD1 *in vitro*, while AF-eTD0 and AF-eTD2 could not bind BmGtsf1L (Fig. 4D).

**Fig 4.**
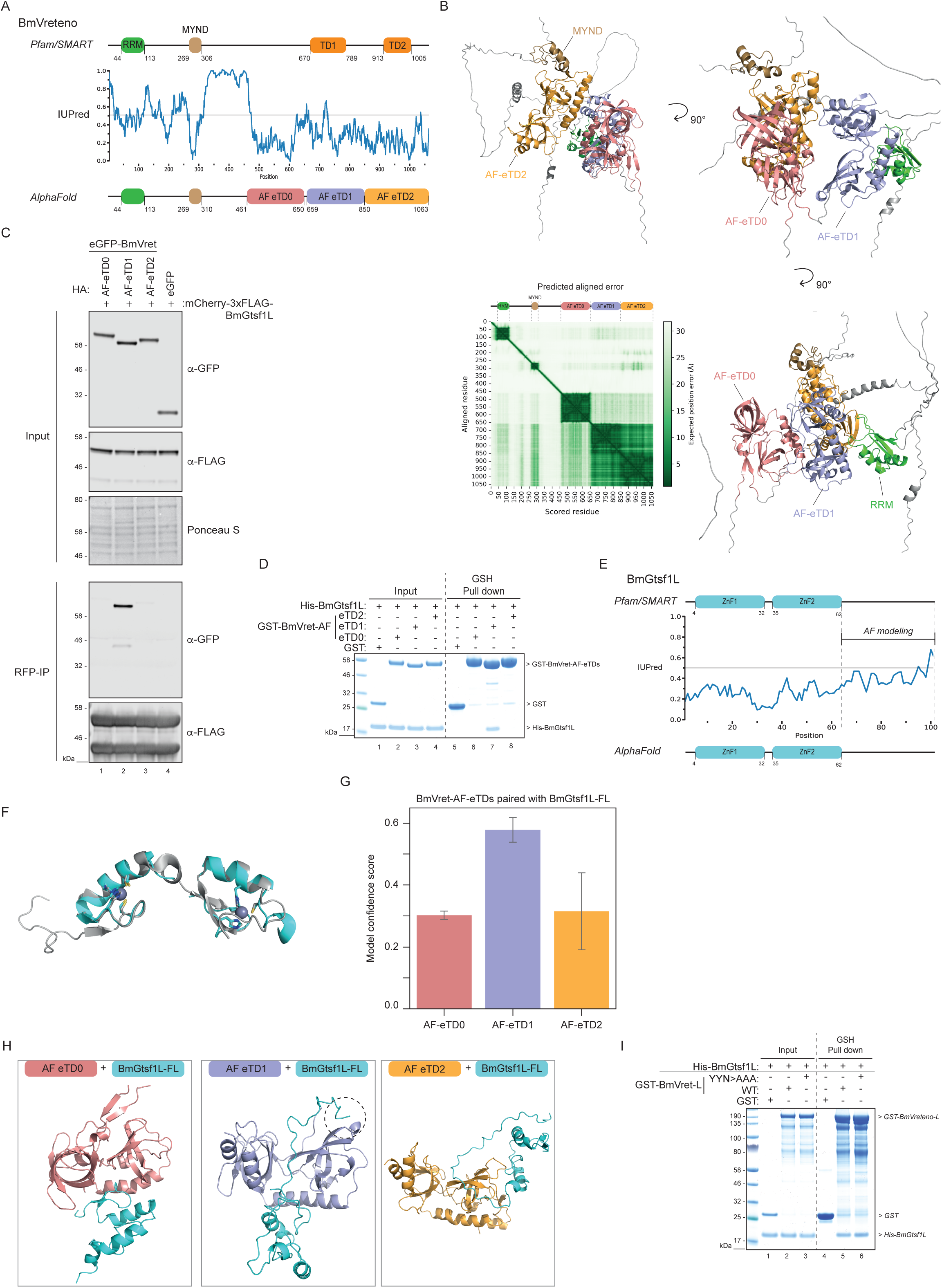
BmVreteno AF-eTD1 interacts with a C-terminal motif in BmGtsf1L. a) Domain organization of BmVreteno based on Pfam/SMART annotations (top) or domain annotations from AlphaFold predictions (bottom). IUPred predictions (center) indicate structural disorder propensities for BmVreteno (values > 0.5 indicate disorder). Disorder scores and amino acid positions are shown on the X-axis and Y-axis, respectively. Abbreviations: RRM=RNA recognition motif; MYND=Myeloid translocation protein 8, Nervy and DEAF-1; TD=Tudor domain; AF-eTD=AlphaFold predicted extended Tudor domain. b) AlphaFold predicted structure of full length BmVreteno shown from different angles. Individual domains within the displayed structure are color coded as in panel (a). BmVreteno domain organization is displayed on top of the PAE matrix. The PAE plot displays the scored residues and aligned residues on the X-axis and Y-axis, respectively. The expected position error in angstroms (Å) is color coded, where dark green color indicates low PAE (high confidence) and white color indicates high PAE (low confidence). c) Transfection of BmN4 cells with mCherry-3xFLAG-BmGtsf1L together with individual eTD domains of BmVreteno, carrying an HA-eGFP tag. Transfection of HA-eGFP served as a control. An RFP immunoprecipitation was performed on BmN4 lysates and input as well as elution samples were resolved by SDS-PAGE. Proteins were detected by Western blot using the indicated antibodies and Ponceau S staining served as a loading control. d) Analysis of the interaction between individual eTDs of BmVreteno and BmGtsf1L by GSH pull-down assays. GST alone or GST-BmVreteno-AF-eTDs were incubated with His-BmGtsf1L. Input and elution fractions were analyzed by SDS-PAGE followed by Coomassie staining. e) Domain organization of BmGtsf1L based on Pfam/SMART annotations (top) or domain annotation from AlphaFold predictions (bottom). IUPred predictions (center) indicate structural disorder propensities for BmGtsf1L. Disorder scores and amino acid positions are shown on the X-axis and Y-axis, respectively. The more disordered C-terminal tail was used for AlphaFold predictions (related to Fig. 5). Abbreviation: ZnF1=zinc finger 1; ZnF2=zinc finger 2. f) Superimposition of the structure of the Zn fingers from MmGtsf1 (in grey, PDB:6X46) with the predicted structure of the Zn fingers of BmGtsf1L by AlphaFold (in cyan). Zinc-binding residues within MmGtsf1 and BmGtsf1L that coordinate zinc ion-binding are displayed as sticks. g) Bar chart showing the different model confidence scores that were obtained from AlphaFold predictions (Y-axis) using individual AF-eTDs of BmVreteno (X-axis) that were paired with full length BmGtsf1L (error bars indicate standard deviation of the five predicted models). h) AlphaFold-predicted structures for each individual eTD of BmVreteno with full length BmGtsf1L. The inset in the middle panel shows that the C-terminal tail of BmGtsf1L establishes contacts with the ordered structure of BmVreteno AF-eTD1. i) Analysis of the interaction between BmVreteno-L full length wildtype (WT) and the aromatic cage mutant (YYN>AAA) with BmGtsf1L by GSH pull-down assays. GST alone or GST-BmVreteno-L was incubated with His-BmGtsf1L. Input and elution fractions were analyzed by SDS-PAGE followed by Coomassie staining.

### BmVreteno AF-eTD1 interacts with a C-terminal motif in BmGtsf1L

Results from the previous sections indicate that a region around the W99 residue in the disordered C-terminal tail of BmGtsf1L can bind to AF-eTD1 pointing to the possibility that this interaction is mediated by a so called short linear motif – folded domain interaction (van Roey *et al*, 2014). Various reports suggest that AlphaFold has some ability to predict domain-motif interfaces between two submitted protein sequences (Akdel *et al*, 2022; Tsaban *et al*, 2022). However, prior to probing AlphaFold for interface prediction between BmVreteno and BmGtsf1L, we first tested whether AlphaFold can predict the structure of full length BmGtsf1L with high confidence. AlphaFold confidently predicted the two N-terminal Zn-finger domains and a disordered C-terminal tail in line with Pfam/SMART domain annotations and IUPred disorder propensity predictions (Fig. 4E, Suppl Data 1). Superimposition of the BmGtsf1L Zn fingers with the resolved structure of mouse Gtsf1 (PDB: 6X46) showed a very similar overall structure (Fig. 4F). Each MmGtsf1 Zn finger coordinates the binding of an individual zinc ion (Ipsaro *et al*, 2021). Displaying the zinc-coordinating residues of BmGtsf1L revealed that AlphaFold accurately modeled these residues, despite the fact that AlphaFold cannot model the zinc ions themselves (Fig. 4F).

Encouraged by these observations we submitted full length BmGtsf1L and full length BmVreteno for interface prediction by AlphaFold. Unfortunately, predicted structural models were of very low model confidence (at most 0.27) and docked the BmGtsf1L Zn finger domains between AF-eTD0 and AF-eTD2 of BmVreteno in an unlikely mode of binding that also contradicts our experimental results (Suppl Data 1).

Next, we submitted sequences of individual AF-eTDs from BmVreteno with the full length sequence of BmGtsf1L for interface prediction. Interestingly, while predictions involving AF-eTD0 and AF-eTD2 resulted again in low confidence predictions, structural models involving AF-eTD1 resulted in substantially higher model confidences (Fig. 4G, Suppl Data 1). Inspection of the structural models revealed that AlphaFold predicted binding of a region involving W99 in BmGtsf1L to AF-eTD1 exclusively (Fig. 4H), in line with our experimental data. Interestingly, AlphaFold docked W99 of BmGtsf1L into a small hydrophobic pocket of AF-eTD1 that was different from the aromatic cage (see below). Indeed, BmGtsf1L can still interact with BmVreteno *in vitro* when the aromatic cage was disrupted (Fig. 4I). This data would suggest that BmAgo3 and BmGtsf1L both bind to BmVreteno AF-eTD1 while using different interaction interfaces on AF-eTD1.

### AF predicts Gtsf1L motif binding to a novel hydrophobic pocket on the BmVreteno AF-eTD1 domain

Despite these encouraging agreements between our experimental data and AlphaFold predictions, the structural models of the interface between AF-eTD1 and full length BmGtsf1L were still only of moderate model confidence (max. 0.64). To further gain in prediction accuracies, we *in silico* fragmented the unstructured C-terminal tail of BmGtsf1L (39 AA in length) starting off with a fragment of five residues in length at the start, middle, and end of the C-terminal tail and gradually extending these fragments by five residues in each step (Fig. 5A). We submitted each fragment individually for interface prediction by AlphaFold with each of the three eTD domains of BmVreteno. This resulted in 72 prediction runs in total (Table S2). Since the C-terminal fragments were overlapping among each other we were able to compute the fraction of prediction runs involving a specific pair of residues from BmVreteno and BmGtsf1L where this pair of residues was predicted to be in contact with each other. This computed fraction was visualized as a heatmap between all residues from BmVreteno AF-eTD domains that were observed to be at least once in contact with a residue from the C-terminal tail of BmGtsf1L and *vice versa* (Fig. 5A). This residue-residue contact heat map revealed a clear hotspot of residues within AF-eTD1 and residues in BmGtsf1L including W99 and residues close by that were consistently predicted to be in contact with each other (Fig. 5A). No such hotspot was observed for AF-eTD0 and AF-eTD2 suggesting that AlphaFold specifically predicted an interface between AF-eTD1 and a motif in BmGtsf1L involving W99. Importantly, model confidences reached 0.87 for these fragment pairs (Table S2).

**Fig 5.**
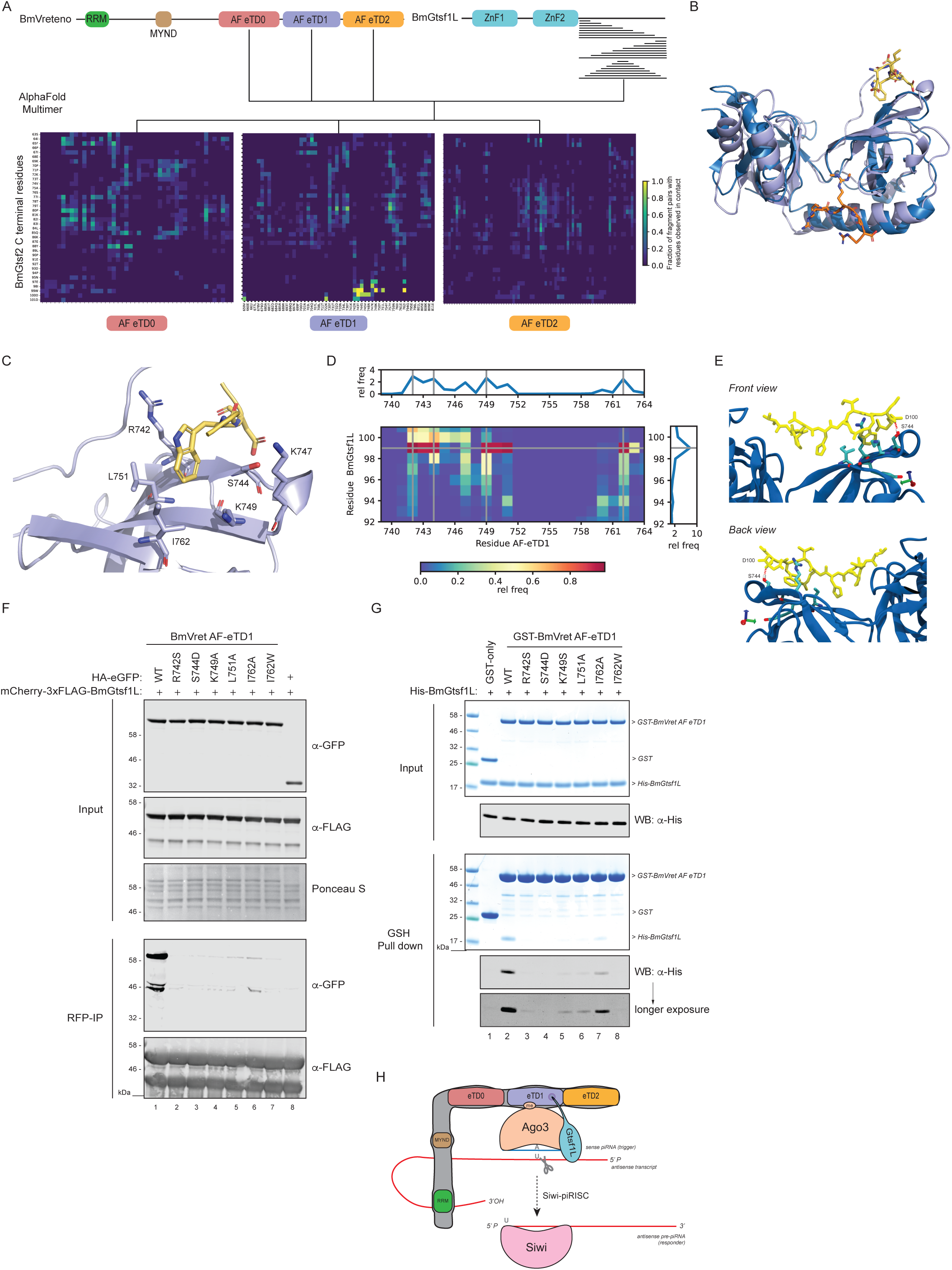
The Gtsf1L motif binds to a novel hydrophobic pocket on the BmVreteno AF-eTD1 domain. a) AlphaFold-based domain organization of BmVreteno and BmGtsf1L and a schematic overview of the fragmentation approach of the C-terminus of BmGtsf1L. BmGtsf1L fragments were paired for interface predictions with AlphaFold with each of the eTDs of BmVreteno (top). The frequency by which a pair of residues, one from BmVreteno and one from BmGtsf1L, was predicted to be in contact with each other among all fragment pairs submitted to AlphaFold that contained this residue pair is visualized as a heatmap for each individual eTD domain of BmVreteno. Only residues of BmVreteno and BmGtsf1L that were at least observed once to be in contact with a partner residue are displayed on the X and Y-axis, respectively. b) Superimposition of eTudor11 from *Drosophila* Tudor (in dark blue, PDB: 3NTH) crystallized with a peptide containing a methylated arginine residue of Aubergine (orange) with the structural model of AF-eTD1 (light blue) and the C-terminal five residue-long peptide of BmGtsf1L (in yellow). Peptide residues are represented as sticks. c) Zoom-in on the novel hydrophobic binding pocket of BmVreteno AF-eTD1 (light blue) and contacts between the hydrophobic residues (shown as sticks) with BmGtsf1L W99 and D100 residues (shown as yellow sticks). d) Contact map of BmVreteno AF-eTD1 with the BmGtsf1L 10-AA residue peptide, predicted by atomistic molecular dynamics simulations. The plot summarizes ten runs of one microsecond each. Blue color in the heatmap indicates a low relative frequency of contacts between the BmGtsf1L-BmVreteno residues and red indicating a high frequency of contacts. Marginal plots that display the relative frequency (rel freq) show the relative probability of a residue interacting with residues from the binding partner, which is the sum of the probability for each column (for sum of the contacts along the AF-eTD1 sequence) or row (for the sum of the contacts along the BmGtsf1L 10-AA residue peptide sequence). e) Snapshot on the novel hydrophobic binding pocket of BmVreteno AF-eTD1 (blue) and contacts between residues R742, S744, K749, and I762 (shown as sticks) with BmGtsf1L C-terminal 10-AA residues (shown as yellow sticks). The snapshot additionally displays the BmVreteno-S744 residue that can form a hydrogen bond with the backbone carbonyl of BmGtsf1L-D100. f) Anti-RFP (BmGtsf1L) immunoprecipitation from BmN4 lysates made from cells that were transfected with HA-eGFP-tagged BmVreteno AF-eTD1 variants. Cells were co-transfected with mCherry-BmGtsf1L. Transfection of HA-eGFP served as a control. Proteins from input and elution samples were resolved by SDS-PAGE, followed by Western blot detection using the indicated antibodies. Ponceau S staining served as a loading control. g) *In vitro* GSH pull-down assay for GST alone or for GST-BmVreteno-AF eTD1 variants incubated with His-tagged BmGtsf1L. Proteins from input and elution fractions are separated by SDS-PAGE and detected by Coomassie staining. For more sensitive detection, a fraction of the same input and elution samples were in parallel subjected gel electrophoresis followed by Western blot detection using anti-His antibodies. h) Model showing that a novel binding interface on BmVreteno AF-eTD1 facilitates the binding of BmGtsf1L (via the hydrophobic pocket) and BmAgo3 (aromatic cage).

We superimposed the structural model involving AF-eTD1 and the last 5 residues of BmGtsf1L (Suppl Data 1) with the solved structure of eTD11 of the *Drosophila* Tudor protein in complex with a synthetic peptide representing the methylated arginine residues of Aubergine (PDB: 3NTH, Fig. 5B) (Liu *et al*, 2010). This clearly shows that the predicted interface between BmVreteno and BmGtsf1L does not involve the aromatic cage and lies on the opposite site of AF-eTD1. Closer inspection of the interface revealed that W99 of BmGtsf1L is predicted to bind in a hydrophobic pocket formed by the side chains of R742, K747, K749, L751, and I762 of AF-eTD1 (Fig. 5C). Furthermore, the conserved D100 of BmGtsf1L is predicted to be in contact with K722 and R742 suggesting charge-charge contacts. This is also true for BmGtsf1L-D101 and BmVreteno-K747 (Fig. 5C). To understand why AlphaFold predictions and experimental results suggest that BmGtsf1L motif-binding is specific to the AF-eTD1 domain of BmVreteno, we superimposed the structural models of all three AF-eTD domains. We observed that the described hydrophobic pocket as well as the aromatic cage are specific to AF-eTD1 (Fig. S4A).

To gain further confidence in the stability of the predicted interface between BmGtsf1L and BmVreteno, we performed atomistic molecular dynamics simulations using the AlphaFold structural model involving AF-eTD1 of BmVreteno and the ten last residues of BmGtsf1L as starting point. In nine out of ten 1 μs simulations we observed that W99 anchors the BmGtsf1L motif into the predicted hydrophobic pocket of AF-eTD1 (Fig. 5D). However, in one of the ten simulations, W99 moves away from the shallow hydrophobic pocket suggesting that additional contacts between both proteins are required to further stabilize the interaction. Our contact analysis of the ten simulation runs demonstrate that the flanking residues I98, D100 and D101 also contribute to anchoring the BmGtsf1L motif, whereas the remaining part of the BmGtsf1L peptide forms fewer contacts with the AF-eTD1 domain and is highly dynamic (Fig. 5D, 5E). On average W99 is interacting with BmVreteno residues R742 and K749 as well as I762. Movie S1, which visualizes one of the ten trajectories, shows how W99 can be ‘sandwiched’ between the side chains of R742 and K749. The simulations also suggested an important contribution of S744 in AF-eTD1 to Gtsf1L motif-binding by mostly interacting with W99 but also with D100 (Fig. 5E). S744 forms transient hydrogen bonds with I98 and D100 of BmGtsf1L (Fig. 5E, S4B & S4C), with the proton of the S744 OH group interacting both with the carboxyl group of the D100 side chain and the D100 backbone carbonyl (Fig. S4C). Interestingly, W99 and D100 also display high conservation scores across orthologous Gtsf sequences (Fig. S1A), suggesting that this may be a conserved mode of binding.

### Experimental verification of the AF-predicted Gtsf1L-BmVreteno interface

We set out to further probe this predicted mode of binding using mutagenesis. To this end, we selected residues within AF-eTD1 that contribute to forming the hydrophobic pocket or/and mediate interaction with the aspartate (D100, D101) residues in the Gtsf1L motif. Individual point mutations (R742S, S744D, K749S, L751A and I762A) were generated for which we hypothesized that these would perturb the formation of the hydrophobic pocket. We also generated a point mutant (I762W) in which the hydrophobic pocket would be filled and as such would sterically hinder the binding of BmGtsf1L. These mutations were also designed such that an overall impact on the stability and folding of AF-eTD1 should be minimal. Immunoprecipitations on BmN4 cell extracts derived from cells that were co-transfected with BmGtsf1L and the panel of AF-eTD1 mutants, showed that the interaction between BmGtsf1L and AF-eTD1 was abolished in all the variants that we tested (Fig. 5F). Using recombinant GST-tagged AF-eTD1 variants in GSH pull-down assays we also observed that the direct interaction between AF-eTD1 and BmGtsf1L was strongly impaired by these mutations (Fig. 5G).

Finally, we studied the effect of these mutations on the BmGtsf1L-BmVreteno interaction in the context of full length proteins. We assessed these interactions through co-IP experiments, as well as via subcellular localization. Mutations within the hydrophobic pocket on AF-eTD1 were introduced into full length BmVreteno and were co-transfected with BmGtsf1L. By microscopy, it was apparent that all the hydrophobic pocket mutants resulted in nuclear BmGtsf1L localization (Fig. S4D). This indicates a loss of binding, since normally BmVreteno overexpression results in BmGtsf1L exclusion from the nucleus. In addition, we find that BmGtsf1L is still present in granules of BmVreteno mutants, which could be explained by the presence of endogenous BmVreteno that can form a complex with the transiently expressed BmVreteno mutant and suffices to recruit a small fraction of BmGtsf1L. Mutation of the aromatic cage did not result in nuclear BmGtsf1L (Fig. S4D), consistent with it not playing a role in BmGtsf1L binding.

Immunoprecipitations of BmGtsf1L revealed that single point mutations within the hydrophobic pocket had little effect on BmVreteno retrieval (Fig. S4E). More significant reduction in binding could be observed for the serine (S744D) and isoleucine (I762W) mutations that also showed the strongest effects in our *in vitro* assay (Fig. S4E, 5E). A stronger loss of interaction was observed when generating double mutants for the hydrophobic pocket residues that initially showed no or marginal effects when compared to wildtype BmVreteno. We note that the full length BmVreteno that we transiently expressed likely still dimerizes or oligomerizes with endogenous BmVreteno, which in turn would still be able to interact with endogenous BmAgo3 (via the aromatic cage). This might lead to the observed residual BmGtsf1L binding. Taken together, our data reveal a novel binding interface on BmVreteno AF-eTD1 that facilitates the simultaneous binding of BmGtsf1L and BmAgo3 (Fig. 5H).

## Discussion

In this study we show that one of the eTudor domains of BmVreteno acts on its own as a molecular scaffold to bring piRNA-loaded BmAgo3 and BmGtsf1L in close proximity. Given that BmGtsf1L is required for efficient cleavage of an RNA target (Arif *et al*, 2022) and that BmVreteno provides an environment that promotes the handover of the cleaved target to empty Siwi (Murakami *et al*, 2021), we propose that the interactions that we identify may help to restrict BmAgo3 cleavage to a molecular surrounding in which its cleavage products can fuel piRNA biogenesis, and to prevent futile BmAgo3 cleavage events. As the targets of BmAgo3 are antisense transcripts, its cleavage activity will not directly contribute to transposon silencing. Only cleavage in the presence of empty Siwi protein will be beneficial to transposon silencing. Therefore, making BmAgo3 cleavage dependent on BmGtsf1L and confining this to the BmVreteno environment would represent an effective way of optimizing BmAgo3 cleavage effectivity.

While it has been revealed that BmVreteno can establish an environment where piRNA-loaded BmAgo3 and empty Siwi are brought together, it is not fully understood how empty Siwi is provided (Nishida *et al*, 2020). It is possible that, in analogy to *Drosophila* Krimper, other eTudor domains of BmVreteno may bind unloaded Siwi (Sato *et al*, 2015; Webster *et al*, 2015). Using AlphaFold modeling, we uncovered in total three eTudor domains in BmVreteno. Two of these do not have an intact aromatic cage, suggesting they may bind empty, unmethylated Siwi.

We and others have shown that BmVreteno is spliced in two isoforms: Long (L) and Short (S). In addition, we know that these two isoforms form heterodimers. *In vitro* RNA cross-linking experiments have revealed that the RRM domain contained within BmVreteno-L binds RNA (Nishida *et al*, 2020). Therefore, it is tempting to speculate that the BmVreteno heterodimer can bring in one target RNA molecule, which is then processed by BmAgo3 and whose 3’-end cleavage product is subsequently loaded onto empty Siwi. Nonetheless, a single BmVreteno-L molecule could in principle also recruit both BmAgo3, Siwi and at least one more protein, so the question why BmVreteno heterodimerizes remains unanswered.

BmVreteno homologs that are expressed within the germ cells of flies (DmVreteno), fish and mice (Tdrd1) are all required for piRNA biogenesis but have different domain organizations (Handler *et al*, 2011; Huang *et al*, 2011; Zamparini *et al*, 2011; Vagin *et al*, 2009; Reuter *et al*, 2009). *Drosophila* and silkworm Vreteno contain an RRM domain and a MYND domain followed by two or three eTudor domains, respectively. Notably, the expression of two Vreteno isoforms seems to be restricted to silkworm. However, in addition to Vreteno, flies also express a Vreteno-like protein in their ovaries, which is called Veneno (Brosh *et al*, 2022). DmVeneno has a very similar domain architecture compared to DmVreteno but is lacking an N-terminal RRM and, as such, mimics the domain organization of the BmVreteno-S isoform (Nishida *et al*, 2020; Brosh *et al*, 2022). Vreteno and Veneno orthologs can also be found in the mosquito species *Aedes aegypti*. Here, Veneno acts as an adaptor protein that brings the ping-pong partners Piwi5 and Ago3 in close proximity for viral piRNA biogenesis (Joosten *et al*, 2019). It would be interesting to study whether Veneno and Vreteno can dimerize and if they co-localize within the nuage of flies and mosquitoes. The mouse and fish Vreteno homologs (Tdrd1) also lack an RRM domain but contain four eTudor domains instead, raising the question how target RNA is provided within these complexes to facilitate *de novo* piRISC assembly. Multivalent interactions within the nuage that are (in part) established by Tudor domains may play an important role here.

The above examples illustrate that many nuage-residing proteins contain multiple eTudor domains, which contribute to the assembly of this phase-separated structure through the formation of multivalent interactions (Chen *et al*, 2011). Importantly, the depletion of a single nuage component can affect nuage integrity and concurs with a significant reduction in piRNA levels. Interestingly, however, in *C.elegans* most eTudor-domain containing proteins that reside in germ granules only harbor one eTudor domain. So how can a single eTudor domain establish a binding platform to recruit multiple proteins? In this study we reveal that a single eTudor domain (AF-eTD1) of BmVreteno can do so by establishing dual binding interfaces. The aromatic cage facilitates the binding of piRNA-loaded, methylated BmAgo3, whereas the hydrophobic pocket allows for binding of BmGtsf1L. The BmGtsf1L C-terminal residues (W99, D100) that are indispensable for binding to the BmVreteno hydrophobic pocket are broadly conserved, indicating that Gtsf proteins might have a preserved mode of binding, which corresponds to a novel type of domain-linear motif interaction. However, more structural studies are needed to understand to which extent the hydrophobic pocket is conserved among other eTudor domains. Interestingly, we recently uncovered another, novel binding interface on an eTudor domain of the *C.elegans* protein TOFU-6 (Podvalnaya *et al*, 2023). This implicates that eTudor domains are much more versatile in establishing multivalent interactions than previously anticipated.

In this study we developed a successful strategy based on AlphaFold for the prediction of protein interaction interfaces involving linear motifs. We note that interface predictions by AlphaFold always return the two protein fragments in contact with each other, making it most of the time very difficult to distinguish good from bad structural models simply by visual inspection. Confidence in reported structural models can be gained from metrics such as the model confidence that is computed by AlphaFold and, as we showed, the recurrent observation of residues predicted in contact with each other when alternating the length of protein fragments submitted for interface prediction. Our work also suggests that interface predictions with AlphaFold using full length proteins might be unsuccessful but more systematic studies are needed to confirm this. Our results further indicate that AlphaFold is able to extrapolate from the training set of protein structures within the PDB to accurately predict protein interaction interfaces it has never seen before. Physics-based models such as molecular dynamics simulations as we employed here also offer a route to investigate and critically assess the importance of binding interfaces predicted by AlphaFold (Zhang *et al*, 2023).

A recent study from Arif *et al*. revealed that Gtsf proteins contribute to the piRNA-guided endonuclease activity of PIWI proteins *in vitro* (Arif *et al*, 2022). The authors proposed a model in which the binding of Gtsf would induce a conformational change in the piRISC-PIWI complex upon pairing with its RNA target. While our manuscript was in preparation another paper reported that BmGtsf1L associates with BmAgo3 and enhances its slicing activity, whereas BmGtsf1 specifically increases Siwi endonuclease activity (Izumi *et al*, 2022). Gtsf’s function to potentiate PIWI slicing is evolutionarily conserved and the strongly evolved C-terminal tail of Gtsf proteins seems to confine binding specificity to its PIWI partner protein (Arif *et al*, 2022). In flies and mouse, conserved aromatic residues within the C-terminus of Gtsf1 contribute to PIWI binding (Dönertas *et al*, 2013; Ohtani *et al*, 2013; Yoshimura *et al*, 2018) and PIWI target cleavage kinetics (Arif *et al*, 2022). However, no direct interaction between the two has thus far been detected *in vitro*. In this study we show that the novel linear motif within the C-terminus of BmGtsf1L is indeed involved in its association with piRNA-loaded BmAgo3, but that this interaction is established through BmVreteno. Therefore, it is tempting to speculate that the association between Gtsf proteins and PIWI proteins in flies and mouse is possibly also mediated via an eTudor domain that thus far has not been uncovered.

To conclude, our studies start to address the question of why PIWI proteins evolved the requirement of a co-factor for target cleavage. Given that other Argonaute proteins can efficiently cleave target RNA without such co-factors, it seems reasonable to pose that the Gtsf dependence of PIWI proteins serves a purpose. We propose that Gtsf proteins are required to dictate where and possibly when target RNAs are cleaved by PIWI proteins to allow for piRNA amplification. Interestingly, in flies and mouse Gtsf proteins also contribute to PIWI-induced transcriptional gene silencing. However, the exact role of Gtsf1 in this process, which does not involve target RNA cleavage, still remains elusive (Dönertas *et al*, 2013; Ohtani *et al*, 2013; Yoshimura *et al*, 2018; de Fazio *et al*, 2011). Perhaps Gtsf proteins restrict conformational changes of PIWI proteins upon target recognition to loci of strong homology, preventing the establishment of transcriptional silencing at erroneous loci. Further studies will be required to test these ideas.

## Materials & Methods

### BmN4 cell culture and transfection

BmN4 cells (a kind gift of Ramesh Pillai) were cultured at 27°C in IPL-41 insect medium (Gibco) supplemented with 10% FBS (Gibco) and 0.5% Pen-Strep (Gibco). 24h prior to transfection, ∼4 × 10^6^ cells were seeded in a 10-cm dish (using one 10-cm dish for each condition). Cells were transfected with plasmid DNA using X-tremeGene HP (Roche) transfection reagent, according to the manufacturer’s instructions. 72h post transfection cells were harvested, washed once in 5 mL ice-cold PBS and once more in 1 mL ice-cold PBS. Subsequently, cells were pelleted by centrifugation for 5 min at 500×*g* at 4°C and frozen at −80°C.

### RNAi in BmN4 cells

For preparation of dsRNA, template DNAs were prepared by PCR using primers that contained flanking T7 promoter sequences. Primers for preparation of dsRNA can be found in Supplementary Materials (Table S3). dsRNA was generated by *in vitro* transcription using the HiScribe T7 kit (NEB), according to the manufacturer’s instructions. Transcribed RNA was purified by phenol/chloroform extraction, precipitated with ethanol and annealed in water. For dsRNA-mediated gene knockdown, ∼2 × 10^6^ BmN4 cells were transfected with 10 µg of dsRNA using X-tremeGene HP. 72h after transfection, cells were again transfected with dsRNA and the dsRNA-treatment was repeatedly performed every 3 days for at least three times for BmGtsf1L depletion and four times for BmVreteno or BmAgo3 knockdown.

### Generation of stable cell lines

For the generation of 3xFLAG-eGFP, 3xFLAG-BmAgo3 and 3xFLAG-Siwi stable cell lines, ∼4 × 10^6^ BmN4 cells were seeded in a 10-cm dish. Cells were transfected with 10 µg of plasmid DNA (Table S3) and cultured under Puromycin (Gibco) selection (5 µg/mL) for at least four additional weeks. Stable integration of plasmid DNA was verified by Western blot. The HA-BmGtsf1L-eGFP stable cell line was generated in a similar manner. All stable cell lines are polyclonal.

### Plasmid construction

For expression of plasmids in BmN4 cells all genes were PCR amplified using BmN4 cDNA and then cloned into the pBEMBL vector (kind git of Ramesh Pillai), which harbors an OpIE2 promoter and an OpIE2 polyA tail (Xiol *et al*, 2012). The plasmids that were used to generate stable cell lines additionally contain a puromycin cassette, where the BmA3 promoter drives the expression of the puromycin-N-acetyltransferase (*pac*) gene, followed by the OpIE2 polyA sequence.

For recombinant protein expression in *E.coli*, coding sequences were cloned into the pET28a(plus) vector that contains an N-terminal (HIS)6-tag or into the pGEX-6p vector for GST-tagged protein expression (kind gift from H. Ullrich lab). All primers, vector backbones and detailed cloning strategies can be found in Supplemental Materials (Table S3).

### Immunoprecipitations

Directly before use, BmN4 cell pellets were thawed on ice and lysed in 1 mL IP-150 Lysis Buffer (30mM Hepes [pH7.4], 150mM KOAc, 2mM Mg(OAc)2 and 0.1% Igepal freshly supplemented with EDTA-free protease inhibitor cocktail and 5mM DTT) for 1h by end-over-end rotation at 4°C. Cells were further lysed by passing the lysate ten times through a 20-gauge syringe needle followed by five passes through a 30-gauge needle. Cell debris was pelleted by centrifugation at 17,000×*g* for 20 min at 4°C. Supernatant fractions were collected and subjected to immunoprecipitations. In case of RNase treatment, 20 µL of RNaseA/T1 (Thermo Scientific, #EN0551) was added to 1 mL of IP-150 lysis buffer prior to lysis (according to the manufacturer’s instructions).

Immunoprecipitations were performed using Pierce™ Anti-HA Magnetic Beads (30 µL bead suspension per reaction, ThermoFischer, #88836), Anti-FLAG M2 Magnetic Beads (20 µL bead suspension per reaction, Sigma, #M8823) or GFP/RFP-Trap Magnetic Agarose beads (15 µL bead suspension per reaction, Chromotek). When using endogenous antibodies, 3 µg of affinity-purified antibodies were coupled to 15 µL of pre-equilibrated Protein G Dynabeads (30 µL bead suspension, Invitrogen) for one reaction. Normal rabbit IgG (Cell Signaling, #2729) or mouse non-immune serum (n.i.) served as controls. Beads and antibodies were incubated for 1h at 4°C by end-over-end rotation in 500 µL of IP-150 Lysis Buffer. Beads-conjugated antibodies were then washed for three times in 1 mL of IP-150 Lysis Buffer. Equilibrated beads were subsequently incubated with the BmN4 cell lysate and incubated overnight by end-over-end rotation at 4°C. The next day, immunoprecipitated complexes were washed five times using 1 mL of IP-150 Lysis Buffer and were subsequently used for immunodetection using Western Blot analysis.

### Western Blot

Samples were prepared in 1x Novex NuPage LDS sample buffer (Invitrogen) supplemented with 100mM DTT and were heated at 95°C for 10 min prior to resolving on a 4-12% Bis-Tris NuPage NOVEX gradient gel (Invitrogen) in 1x Novex NuPAGE MOPS SDS Running Buffer (Invitrogen) at 140V. For the detection of endogenous BmGtsf1L, proteins were resolved on a 15% Bis-Tris polyacrylamide gel. Separated proteins were transferred to a nitrocellulose membrane (Amersham) overnight at 20V using 1x NuPAGE Transfer Buffer (Invitrogen) supplemented with 10% methanol. The next day, the membrane was blocked for 1h in 1x PBS-Tween (0.05%) supplemented with 5% skim milk and incubated for 1h with primary antibodies diluted in blocking buffer (1:1,000 anti-Flag; 1:1,000 anti-GFP; 1:1,000 anti-HA; 1:1,000 anti-actin, 1:2,500 anti-tubulin; 1:1,000 for all endogenous antibodies). Subsequently, the membrane was washed three times for 5 min in PBS-Tween, prior to 1h incubation with the secondary antibody, using 1:10,000 IRDye 800CW Goat anti-mouse and IRDye 680LT Donkey anti-rabbit IgG (LI-COR) and imaged on an Odyssey CLx imaging system (LI-COR). Secondary antibodies used for chemiluminescence-based detection were 1:1,000 rat monoclonal anti-mouse Ig HRP (Clone eB144, Mouse TrueBlot ULTRA, Rockland #18-8817-30), 1:10,000 goat anti-rabbit IgG HRP-linked antibody (Cell Signaling Technology, #7074), 1:10,000 horse anti-mouse IgG, HRP (Cell Signaling Technology, #7074). Chemiluminescence signals were detected using ECL select Western Blotting detection reagent (Cytvia, #GERPN2235) and imaged on a Fusion FX imaging system (Vilber).

### Recombinant protein purification

GST alone as well as GST-3C-BmVreteno variants and fragments were expressed from pGEX6p vectors. His6-thrombin-BmGtsf1L variants were expressed from pET vectors. Transformed plasmids were expressed in *E. coli* (BL21 DE3 codon+, Agilent) overnight at 18°C using 0.5mM IPTG in LB media. Cells were lysed in ice-cold lysis buffer (30mM Tris-Cl pH 8.0, 500mM NaCl, 0.5mM TCEP, 5% glycerol, EDTA-free cOmplete protease inhibitor cocktail and additional 10mM imidazole pH 8.0 for BmGtsf1L purifications), using a CF1 continuous flow cell disruptor from constant systems at 29 kpsi and cleared by centrifugation at 40,000×g for 30 min at 4°C. Recombinant proteins were affinity-purified from cleared lysates using a NGC Quest Plus FPLC system (Biorad) and GSTrap HP (GST-tagged BmVreteno and GST), or HisTrap HP (His6-tagged BmGtsf1L) 5 mL columns (Cytiva), according to the manufacturers protocols. Eluted proteins were concentrated using Amicon spin concentrators (Merck Millipore) and subjected to gel filtration (Superdex 75 and 200 16/60 pg, Cytiva, in 25mM Na-Hepes, 300mM NaCl, 10% Glycerol, pH 7.4).

To obtain an untagged BmVreteno (181-386) antigen fragment for immunization, the GST-tagged fragment from the affinity step was digested with 3C protease (1:100 w/w) overnight at 4°C during dialysis (30mM Tris-Cl pH 8.0, 500mM NaCl, 1mM DTT, 5% glycerol). The digested protein was re-applied to a GSTrap HP 5 mL column to absorb the free GST. The flow through from this step, containing the untagged BmVreteno (181-386) antigen was concentrated using Amicon spin concentrators and subjected to gel filtration (Superdex 75 16/60 pg in PBS). Another round of free GST absorption via GSTrap, followed by gel filtration (Superdex 75 16/60 pg in PBS) was performed to remove residual free GST from the untagged BmVreteno (181-386) antigen.

For all recombinant proteins, peak fractions after the final gel filtration were pooled and protein concentration was determined by using absorbance spectroscopy and the respective extinction coefficient at 280 nm, before aliquots were flash frozen in liquid nitrogen and stored at −80°C.

### GSH pull-downs

Glutathione Sepharose 4B beads (Cityva) were equilibrated (20 µL beads suspension for each reaction) by three washes in PBS containing 0.1% Triton-X100 (PBS-T) and the resin was pelleted by mild centrifugation at 1,000×*g* for 2 min at 4°C. Next, 5µM of GST-BmVreteno was added to the beads together with 10µM of His-BmGtsf1L and samples were incubated for 2h by end-over-end rotation at 4°C. Beads were pelleted by centrifugation at 1,000×*g* for 2 min at 4°C and washed three times with PBS-T. Finally, pelleted beads were resuspended in 25 µL 1x Novex NuPage LDS sample buffer (Invitrogen) supplemented with 100mM DTT and were heated at 95°C for 5 min prior to resolving (40% of the sample) on a 4-12% Bis-Tris NuPage NOVEX gradient gel (Invitrogen) in 1x Novex NuPAGE MES SDS Running Buffer (Invitrogen) at 180V. Proteins on the gel were visualized by staining with InstantBlue Coomassie protein stain (Abcam).

### Antibodies

Monoclonal antibodies for detection and/or immunoprecipitation of endogenous Siwi, BmAgo3, BmSpn-E and BmQin were a kind gift from Mikiko Siomi (Nishida *et al*, 2015). Rabbit polyclonal antibodies for BmAgo3 detection were provided by Ramesh Pillai (Xiol *et al*, 2012). The monoclonal anti-BmGtsf1L antibody was generated in the Siomi lab by immunizing mice with purified GST-tagged full length BmGtsf1L. Fusing myeloma generated hybridomas as described previously (Nishida *et al*, 2015).

The rabbit polyclonal anti-BmVreteno antibody was generated by immunizing rabbits with the affinity purified BmVreteno (186-381) antigen (Eurogentec). 2 mL sulfolink resin (Thermo Fisher Scientific) was covalently conjugated with 3 mg GST-tagged BmVreteno (181-386) according to the manufacturers protocol. 10 mL final bleed of each rabbit serum was incubated with 1 mL GST-BmVreteno (181-386)-conjugated sulfolink resin at 4°C overnight while rotating. After incubation, the resin was washed with PBS containing 0.1% Triton X-100, followed by a wash with PBS in a gravity-flow poly-prep column (Biorad). Elution of polyclonal antibody species was performed using low pH (100mM Glycine-Cl, 150mM NaCl, pH 2.3), followed by immediate neutralization of elution fractions with Tris-Cl pH 8.0. The eluted antibodies were re-buffered using a PD-10 column (PBS, 10% glycerol, 0.05% NaN3) and concentrated to 1 mg/mL using Amicon spin concentrators, before flash freezing in liquid nitrogen and storage at −80°C.

Monoclonal anti-HA was produced in house (clone 12CA5, Core Facility Protein Production). Rabbit polyclonal Anti-HA (Sigma-Aldrich, #SAB4300603), mouse monoclonal anti-Flag M2 (Sigma-Aldrich, #F3165), rabbit polyclonal anti-FLAG (Milipore, #F7425), rabbit polyclonal anti-actin (Sigma-Aldrich, #A5060), mouse monoclonal anti-alpha Tubulin (clone B-5-1-2, Sigma-Aldrich, #T6074), rabbit polyclonal anti-GFP (Origene, #TP401) and mouse monoclonal anti-GFP (clone B-2, Santa Cruz, #sc-9996) are all commercially available.

### Sequence alignment

Clustal W (Larkin *et al*, 2007) and Jalview software (Waterhouse *et al*, 2009) was used for protein alignment and visualization.

### Microscopy

For co-localization studies, approximately 2 × 10^4^ cells were seeded per well in 8-well µ-slide (Ibidi, #80826). The next day, cells were transfected with 100 ng of each corresponding plasmid using X-tremeGene HP. 24h post transfection, live cells were imaged using the Leica TCS SP5 with a 60x oil immersion objective lens. Images were processed using FIJI (Schindelin *et al*, 2012) and Adobe Illustrator software.

### RNA isolation and small RNA sequencing

Per condition, one well of a 6-well plate was seeded with 6 × 10^5^ BmN4 cells 24h prior to transfection with either HA-eGFP, HA-BmAgo3, HA-Siwi or HA-BmGtsf1L using X-tremeGene HP transfection reagent. 72h post transfection, cells were harvested and an anti-HA immunoprecipitation was performed as described above, the experiment was performed in duplicate. Immunopurified RNAs were extracted from beads by adding 1 mL Trizol LS (Invitrogen #10296028), according to the manufacturer’s instructions. The lysate was incubated at RT for 5 min to allow complete dissociation of the nucleoprotein complex. Next, 200 µL of chloroform was added to 1 mL of lysate followed by harsh mixing and centrifugation at 12,000×*g* for 15 min at 4°C. Another round of chloroform extraction was performed and the aqueous phase was transferred to a fresh tube to which 1 volume (500 µL) of ice-cold isopropanol was added for RNA precipitation. RNA pellets were washed twice in 1 mL of 70% ice-cold ethanol and centrifuged at 7,500×*g* for 10 min at 4°C. The RNA pellet was air-dried and dissolved in nuclease-free water.

NGS library prep was performed with NEXTflex Small RNA-Seq Kit V3 following Step A to Step G of Bioo Scientific’s standard protocol (V16.06) using the NEXTFlex 3’ SR Adaptor and 5’ SR Adaptor (5’rApp/NNNNTGGAATTCTCGGGTGCCAAGG/3ddC/and5’ GUUCAGAGUUCUACAGUCCGACGAUCNNNN, respectively). Libraries were prepared with a starting amount of 7 ng and amplified in 25 PCR cycles.

Amplified libraries were purified by running an 8% TBE gel and size-selected for 15 – 35nt. Libraries were profiled in a High Sensitivity DNA Chip on a 2100 Bioanalyzer (Agilent technologies) and quantified using the Qubit dsDNA HS Assay Kit, in a Qubit 2.0 Fluorometer (Life technologies).

All samples were pooled in equimolar ratio and sequenced on 1 Highoutput NextSeq 500/550 Flowcell, SR for 1x 84 cycles plus 7 cycles for the index read.

### Bioinformatic analyses

The quality of raw sequenced reads was accessed with FastQC, Illumina adapters were then removed with cutadapt (-O 5 -m 28 -M 45), reads with low-quality calls were filtered out with fastq quality_filter (-q 20 -p 100 -Q 33). Using information from unique molecule identifiers (UMIs) added during library preparation, reads with the same sequence (including UMIs) were collapsed to remove putative PCR duplicates using a custom script. Prior to mapping, UMIs were trimmed (seqtk trimfq -b 4 -e 4) and library quality re-assessed with FastQC. Reads were aligned against the silkworm (*Bombyx mori*) genome assembly obtained from lepbase GCA_000151625.1 with bowtie v1.1.1 (-l 40 -n 2 -e 70 -m 1 –tryhard –best –strata –chunkmbs 256 –phred33-quals). The locations of repeat elements were also downloaded from lepbase, repeat masker scaffolds (ASM15162v1), converted to genomic location with rmsk2bed. These locations were used to select reads mapping to repeats by intersecting with bedtools intersect (-wa -wb -bed -f 1.0 -nonamecheck) with either the flags -s or -S to determine which small RNAs map sense or antisense, respectively, to the annotated repeats. After filtering, length profiles were obtained by summarizing the length of these reads. Sense/antisense bias was determined by calculating the ratio of reads mapping in the same or the opposite strand for each annotated repeat - repeats with 10 or fewer mapped reads were excluded. Nucleotide bias of piRNAs was determined by summarizing the number of times a base is present in any given piRNA (read sequence) position.

### Mass-spectrometry

About 4 × 10^6^ BmN4 cells were transfected with HA-tagged BmGtsf1L or with HA-eGFP, which served as a control to detect non-specific binders. Cells were harvested 72h post transfection and an anti-HA immunoprecipitation was performed (as described above) on 4 mg total protein lysate. The experiment was performed using two technical duplicates to perform quantitative mass-spectrometry based detection of unique peptides using stable dimethyl isotope labeling (Hsu *et al*, 2003).

#### Protein in-gel digestion

Proteins were separated briefly in a 10% NuPAGE Bis-Tris gel, stained with Coomassie blue and cut into small gel cubes, followed by destaining in 50% ethanol/25mM ammonium bicarbonate. Afterwards, proteins were reduced in 10mM DTT at 56°C and alkylated by 50mM iodoacetamide in the dark at room temperature. Enzymatic digestion of proteins was performed using trypsin (1 µg per sample) in 50mM TEAB (triethylammonium bicarbonate) overnight at 37°C. Following peptide extraction sequentially using 30% and 100% acetonitrile, the sample volume was reduced in a centrifugal evaporator to remove residual acetonitrile. The sample volume was filled up to 100 µL by addition of 100mM TEAB.

#### Dimethyl-labelling

Dimethyl-labelling was performed as previously reported (Boersema *et al*, 2009). Briefly, the digested samples were labelled as “Light” or “Heavy” by adding formaldehyde or formaldehyde-d2, respectively. This was followed by addition of NaBH3CN. Thereafter, the samples were incubated at room temperature with orbital shaking for 1 h. The labelling reaction was quenched by adding ammonia solution. Next, peptides were acidified with formic acid to reach pH ∼3. The paired labelled samples were then combined. The resultant peptide solution was purified by solid phase extraction in C18 StageTips (Rappsilber *et al*, 2003).

#### Liquid chromatography tandem mass spectrometry

Peptides were separated in an in-house packed 30-cm analytical column (inner diameter: 75 μm; ReproSil-Pur 120 C18-AQ 1.9-μm beads, Dr. Maisch GmbH; heated at 40°C) by online reverse phase chromatography through a 105-min non-linear gradient of 1.6-32% acetonitrile with 0.1% formic acid at a nanoflow rate of 225 nL/min. The eluted peptides were sprayed directly by electrospray ionization into a Q Exactive Plus Orbitrap mass spectrometer (Thermo Scientific). Mass spectrometry measurement was conducted in data-dependent acquisition mode using a top10 method with one full scan (mass range: 300 to 1,650 m/z; resolution: 70,000, target value: 3 × 10^6^, maximum injection time: 20 ms) followed by 10 fragmentation scans via higher energy collision dissociation (HCD; normalised collision energy: 25%, resolution: 17,500, target value: 1 × 10^5^, maximum injection time: 120 ms, isolation window: 1.8 m/z). Precursor ions of unassigned or +1 charge state were rejected. Additionally, precursor ions already isolated for fragmentation were dynamically excluded for 20 s.

#### Mass spectrometry data processing and statistical analysis

Raw data files were processed by MaxQuant software package (version 1.5.2.8) (Cox & Mann, 2008) using its built-in Andromeda search engine (Cox *et al*, 2011) and default settings. Spectral data were searched against a target-decoy database consisting of the forward and reverse sequences of the bait proteins (HA-eGFP and HA-BmGtsf1L), *Bombyx mori* proteomes (UniProt 18,382 entries; NCBI 29,282 entries) downloaded on 8th January 2018, a collection of self-cloned *Bombyx mori* genes (28 entries) and a list of 245 common contaminants. Corresponding labels were selected for “Light” (DimethLys0 and DimethNter0) and “Heavy” (DimethLys4 and DimethNter4) labels. A maximum of 3 labelled amino acids per peptide were considered. Trypsin/P specificity was assigned. Carbamidomethylation of cysteine was set as fixed modification. Oxidation of methionine and acetylation of the protein N-terminus were chosen as variable modifications. A maximum of 2 missed cleavages were tolerated. The minimum peptide length was set to be 7 amino acids. False discovery rate (FDR) was set to 1% for both peptide and protein identifications.

For protein quantification, minimum ratio count of two was required. Both the unique and razor peptides were used for quantification. The “re-quantify” function was switched on. The “advanced ratio estimation” option was also chosen. Downstream data analysis was performed in R statistical environment. Reverse hits, potential contaminants and protein groups “only identified by site” were filtered out. Protein groups with at least two peptides including at least one unique peptide were retained.

### AlphaFold predictions

We used the following sequences for AlphaFold predictions:

BmGtsf1L: MDDPFVSCPYNPIHRVPRSRLQRHIVKCEWINPTMIACPYNATHRYTQED MKFHVLNCPSKTSIFPIEKPPKTVASITTPKIILQKEYLPETDPNHEIWDD

BmVreteno:

MSNHSRPQRRREWDPMRDDFNEHTYDVQYADDNAGEQVQLDHTKLYIINI PRGLSEDGIRAAFSKHGKVLSARLSKNPNKRFAIVQFETASEAKLAMMKM NGSEPLNLKISIAHKTIRKTQHDNKDRNYSTSRNGHCSRDEASSISSKGW NMRNLDDVMNNDEIDEIDDMIHEDHDDNLDLELDMLTLKQLKIKEEQLMC KRRLLLRHAEKRQVAPHSSAGRSVLPDGRIVVRNNANETDSAEVEPSFAG AGSESLKTPGLERNASRQCVKCGAPADWYCSRCAITPYCSQTCQTRDWTE RHKSVCHYLAPLKTAGGFEAEATSSKSVSNSTPMRSSHSPPTKQQRGEAD ETDNKAKNIQEPRQNYHRPSNSGPNKNIPGKNQDPRRPATSREAIEEETE ERGARNPKPAEATKDKHHPMNPVTFQRRQLKSNPVVDAQPAPREQQQPAA TRAPEASPTEQRESTRRTLVPDRCLIDSLSEGDVVLVSVELKASECCTKQ GGYVCLSMHEKYESDYQKLCEDYVLDCEADSDEYKIITGDTFSYLSPEDG GWYRARALNTTMAALLDGSKVVYLRMNDKVKKLPAKYSGIPEFCCVLNAD VEVGLNLKCSLLSKTPNGFKVTLENVETEANVGEGEITRWIPEVDYPPPV KNVPVQRSVEIPEVPRPEIKNKSRVILVDATDVQRVFVRPADTRSQKAFD NILQDVLLYGTTAEPLKEPPSKGQTVVSKYTDNLHYRALCKRTSVNKNKY LLEYIEYGNIEITQLNRLYPCPEHLSVTSLASLTSHVQLDTTVGELTPRA LEYIETIKEEEMILTLSSGGDTAQSGAALVNLTLVKNNDNVNKRIEELCT PEWKKLELKGVDVIETERLMYGTALDYIELPAAPFDLQVLDEVGLDSGNI SGCPTNSDYVRYVMTKLPARMREYCESEFGRQPYLPAAEELCIAQLPPSS EWHRAVVLEQILGPGGGTARVLFVDHGNVAEVPVSSLRKMLAEFVTDLPA VACQIVIEDFPKQATAEMLAKARRFMSGPDKARAAQLPVRGCDKQDVGIY AIRVPELLEAMTE

We ran AlphaFold v2.2 (Jumper *et al*, 2021) for all monomeric protein predictions and AlphaFold-Multimer v2.2 (Evans *et al*, 2022) for all protein complex predictions with the following parameters:

--max_template_date=2020-05-14

--db_preset=full_dbs

--use_gpu_relax=False

For every AlphaFold run, 5 models were predicted with one seed per model by setting the following parameter:

--num_multimer_predictions_per_model=1

Out of the five models generated, we used only the model ranked_0 for further processing and interpretation. We defined two residues, one from each protein fragment, to be in contact with each other in predicted AlphaFold models, if at least one heavy atom from one residue is less than 5Å away from any heavy atom from the other residue. Distance measurements between heavy atoms were obtained using the function cmd.distance from PyMOL. The model confidence was extracted from the ranking_debug json file. The PAE matrix was extracted from the pickle file of the model. We used the software, PyMOL (TM) Molecular Graphics System, Version 2.5.0. Copyright (c) Schrodinger, LLC., for the visualization and superimposition of AlphaFold models. The superimposition of the structural model involving AF-eTD1 and the last 5 residues of BmGtsf1L with the solved structure 3NTH was done using the cealign command where AF-eTD1 was set as the mobile entity and the chain A of 3NTH as the target entity for superimposition. For the superimposition of AlphaFold-predicted BmGtsf1L with the solved structure 6X46, we extracted the two Zn fingers from AlphaFold-predicted BmGtsf1L (residue 8 to 34 and 35 to 64 for the two Zn fingers, respectively) and aligned them to the two Zn fingers from chain A of 6X46 (residue 14 to 41 and 48 to 75 for the two Zn fingers, respectively). The align command was used for the superimposition where AlphaFold-predicted BmGtsf1L Zn fingers were set as the mobile entities and the Zn fingers from the first ensemble state of 6X46 as the target entities.

IUPred predictions were obtained by submitting full length sequences to the webserver of IUPred2A (Mészáros *et al*, 2018) and selecting the option IUPred2 long disorder (default) for disorder propensity predictions.

We used the Python libraries, pandas (McKinney, 2010) for data analysis, and Matplotlib (Hunter, 2007) and seaborn (Waskom, 2021) for data visualization.

### Molecular dynamics simulations

We ran atomistic molecular dynamics simulations using the AlphaFold structural model involving AF-eTD1 of BmVreteno and the ten last residues of BmGtsf1L. We used the Amber99SB*-ILDN-q protein force field (Best & Hummer, 2009; Hornak *et al*, 2006; Lindorff-Larsen *et al*, 2010; Best *et al*, 2012) and the TIP4P-D water model (Piana *et al*, 2015). Molecular dynamics simulations were run in GROMACS 2021 (www.gromacs.org) (Abraham *et al*, 2015).

The protein-peptide complex was simulated in a rhombic dodecahedron, with a minimum distance of 12 Å between protein atoms and box edges. 150 mM NaCl were added to the solvated simulation system. The system was energy minimized and equilibrated for 1 ns in using the Berendsen thermostat and barostat at 300 K and 1 bar (Berendsen *et al*, 1984).

We run ten independent simulations starting from the equilibrated starting structure, each with a different set of initial velocities. Each of the ten simulations was run for 1 μs. The Bussi-Donadio-Parinello thermostat was used to maintain a simulation temperature of 300 K (Bussi *et al*, 2007). Parrinello-Rahman barostat was employed to keep pressure at 1 bar (Parrinello & Rahman, 1981). Electrostatics were described by the particle mesh Ewald method (PME). The cut-off for van der Waals interactions was 12 Å.

Simulations were analyzed with the MDAnalysis Python library (Michaud-Agrawal *et al*, 2011; Gowers *et al*, 2016). Two residues were deemed to be in contact if one pair of atoms was within 4.5 Å. The contacts maps from the ten simulation runs, which were started from the same starting structured were averaged to produce a single contact map. Hydrogen bonds were quantified as described by Smith et al (Smith *et al*, 2019).

## Data and material availability

The datasets produced in this study are available in the following databases:

- The accession number for the smRNA-seq data generated in this study is PRJNA940809 (https://www.ncbi.nlm.nih.gov/bioproject/PRJNA940809).
- Mass spectrometry proteomics data: ftp://MSV000091404@massive.ucsd.edu
- All plasmids and reagents are available upon request.

## Acknowledgements

We thank all members of the Ketting lab for fruitful discussions. We would like to thank Ramesh Pillai for providing plasmids, antibodies and the BmN4 cell line. We are very grateful to the Siomi lab for their generous supply of antibodies. The IMB Core Facility Microscopy is acknowledged for support with confocal microscopy. Support by IMB Proteomics Core Facility is gratefully acknowledged (instrument funded by DFG INST 247/766-1 FUGG). In particular, we wish to thank Mario Dejung, Proteomics Core Facility, for his continuous support and for extensive bioinformatic analysis. Support by the IMB Genomics Core Facility and the use of its NextSeq500 (funded by the DFG – INST 247/870-1 FUGG) is gratefully acknowledged. Support from IMB’s IT department and especially help from Christian Diedrich for a local installation of AlphaFold is gratefully acknowledged. The GPU cluster on which all AlphaFold predictions were performed was funded by the Ministry of Science and Health (MWG), Rhineland Palatinate (funding id: TB-Nr.:3658/19).

AWB was supported by the Peter und Traudl Engelhorn Foundation, by the Klaus Tschira Boost Fund and the German Scholars Organization (GSO/KT 14) as well as by an SPP-1935 start-up fund from the Deutsche Forschungsgemeinschaft (DFG, German Research Foundation). MC Siomi is supported by MEXT KAKENHI Grant Number JP19H05466. This work was further supported by the DFG-funded project LU 2568/1-1 granted to KL. L.S.S. acknowledges support by ReALity (Resilience, Adaptation and Longevity), M^3^ODEL and Forschungsinitiative des Landes Rheinland-Pfalz. Further, we gratefully acknowledge the advisory services offered and the computing time granted on the supercomputers Mogon II at Johannes Gutenberg University Mainz, which is a member of the AHRP (Alliance for High Performance Computing in Rhineland Palatinate) and the Gauss Alliance e.V.

## Author contributions

A.W.B, L.S., K.L, and R.F.K. conceived the study and designed experiments. A.W.B. executed all wet lab experiments and performed data analysis. S.S. assisted in wet lab experiments. M.M.M and S.R. performed the protein purifications. T.S and M.C.S generated the anti-BmGtsf1L antibody. C.Y.L and K.L conducted all AlphaFold-related work. L.S. performed atomistic molecular dynamics simulations. A.M.d.J.D. performed small-RNA-seq analysis. A.W.B and R.F.K. supervised the project. A.W.B., L.S., K.L. and R.F.K. wrote the manuscript with input from all authors.

## Supplementary Figure legends

**Fig S1 | BmGtsf1L and BmVreteno both interact with BmAgo3**

a) Domain organization (top) and ClustalW alignment of GTSF proteins from different species. The alignment and conservation scores are depicted using the Jalview software. Residues that are highlighted in blue reveal a 20% identity threshold

b) Western blot detection using the mouse monoclonal anti-BmGtsf1L antibody on naïve BmN4 cell extracts or on BmN4 cells that were transfected with FLAG-BmGtsf1L.

c) Validation of stable integration of FLAG-Siwi, FLAG-BmAgo3 or FLAG-eGFP expression cassettes into BmN4 cells after extensive puromycin selection by Western blot using the indicated antibodies. Anti-actin probing served as a loading control.

d) Control or GFP (BmGtsf1L) immunoprecipitation on BmN4 cell extracts from FLAG-PIWI stable cells that were co-transfected with BmGtsf1L-eGFP. Western blot was performed using the indicated antibodies and anti-actin probing as well as Ponceau S staining served as loading controls.

e) Nucleotide composition of small RNAs that were sequenced from input samples or from anti-HA immunoprecipitated samples.

f) Pfam/SMART-based domain organization of the BmVreteno-Long and BmVreteno-Short isoforms, showing the RNA-recognition motif (RRM), Myeloid translocation protein 8, Nervy and DEAF-1 (MYND) domain, and two C-terminal Tudor domains (TD).

g) Western blot using the rabbit polyclonal anti-BmVreteno antibody on cell extracts from BmN4 cells that were either untransfected or transfected with HA-BmVreteno (L)-FL. Anti-tubulin probing served as a loading control.

h) Validation of anti-BmVreteno antibody specificity on cell extracts from BmN4 cells that were transfected four consecutive times with dsRNA against Luciferase (Luc) or against BmVreteno.

i) IgG or anti-BmVreteno immunoprecipitation on naïve BmN4 cells, followed by Western blot detection of endogenous BmVreteno and BmAgo3.

j) Reciprocal IP on BmN4 cell extracts using non-immune (n.i.) serum or anti-BmAgo3 antibodies as well as endogenous BmVreteno antibodies for Western blot detection of retrieved proteins.

**Fig S2 | BmAgo3 interacts with BmVreteno eTD1 via methylated arginine residues**

a) Multiple sequence alignment of Tudor domains expressed in BmVreteno and its orthologue in *Drosophila* (DmVret). In addition, eTudor11 from the *Drosophila* Tudor protein as well as the eTudor domain of *Drosophila* Tudor-SN (p100) for which crystal structures have been resolved (PDB: 3NTH and 2WAC, respectively) were included. Alignments were performed using Clustal Omega and were processed using Jalview software. Aromatic cage residues are depicted as green boxes and the asparagine residue that is involved in directly binding to the methylated arginine residue (sDMA) is highlighted in yellow. Identical residues (*), conserved substitutions (:) or substitutions by weakly similar residues (.) are indicated below the alignment.

b) Anti-FLAG immunoprecipitation of FLAG-BmAgo3 variants that were transiently expressed in the HA-BmGtsf1L-eGFP stable BmN4 cell line. Transfection of 3xFLAG-mCherry served as a control. Immunoprecipitated proteins were analyzed by Western blot using the indicated antibodies, whereas Ponceau S staining served as a loading control.

**Fig S3 | The BmGtsf1L C-terminus establishes an interaction with BmVreteno**

a) Outlined strategy for the purification of recombinant GST-tagged BmVreteno-S, showing the profiles from the size-exclusion column (left) and the peak fractions that were analyzed by SDS-PAGE followed by Coomassie staining (right).

b) Similar to panel (a), but for GST-BmVreteno-L.

c) Size-exclusion profiles of BmVreteno-S (left) and BmVreteno-L (right) to compare the shift in molecular weight before (green line) and after (blue line) 3C-mediated cleavage of the GST-tag.

d) Pfam/SMART-based domain organization of the BmVreteno-Long isoform, showing the RNA-recognition motif (RRM), Myeloid translocation protein 8, Nervy and DEAF-1 (MYND) domain, and two C-terminal Tudor domains (TD). In addition to the full length (FL) BmVreteno, the C-and N-terminal truncation variants are depicted that are used in panel (e) to analyze the interaction between BmGtsf1L and BmVreteno variants.

e) Co-transfection of BmGtsf1L-eGFP with HA-BmVreteno truncation variants in which different domains were omitted. Transfection of HA-LacZ served as a control. BmGtsf1L was retrieved by GFP-IP and input and elution fractions were analyzed by SDS-PAGE, followed by Western blot using the indicated antibodies. Anti-actin immunodetection served as a loading control.

**Fig S4 | The hydrophobic binding pocket is unique to BmVreteno AF-eTD1 and facilitates BmGtsf1L binding.**

a) Superimposition of the AlphaFold structural models of all three BmVreteno AF-eTD domains. The central inset (closed circle) shows the side view of the aromatic cage, which is only present in AF eTD1 and indicated with a dashed circle. The top inset (closed circle) shows a top view of the novel interface, which is unique to AF eTD1. The hydrophobic pocket is indicated with a dashed circle and the enlarged view additionally shows the docked C-terminal motif of BmGtsf1L.

b) Snapshot on the novel hydrophobic binding pocket of BmVreteno AF-eTD1 (blue) and contacts between the residues R742, S744, K749, and I762 (shown as sticks) with BmGtsf1L C-terminal 10-AA residues (shown as yellow sticks). This snapshot displays the BmVreteno S744 residue that forms a hydrogen bond with the backbone carbonyl of BmGtsf1L I98.

c) Plots showing the distances between the atoms forming the four most important inter-chain hydrogen bonds of the side chain of BmVreteno S744 in two out of the ten 1 μs simulation runs. The run presented in the upper panel displays the hydrogen bond between the side chain of S744 and the backbone carbonyl of BmGtsf1L I98. The simulation presented in the bottom panel reveals that S744 is engaged in different interactions with BmGtsf1L residues I98 and D100. Overall, the simulations reveal that some of the hydrogen bonds are transiently formed and broken.

d) Single-plane confocal micrographs of BmN4 cells co-transfected with different eGFP-BmVreteno constructs (upper panel) and BmGtsf1L-mCherry (middle panel). Yellow triangles indicate a formed granule. Scale bars – 4 µm.

e) Transfection of BmN4 cells with BmGtsf1L-eGFP together with HA-BmVreteno-FL. Transfection of HA-LacZ served as a control. A GFP (BmGtsf1L) immunoprecipitation was performed on BmN4 lysates and input as well as elution samples were resolved by SDS-PAGE. Proteins were detected by Western blot using the indicated antibodies and anti-actin probing as well as Ponceau S staining served as a loading control.

