## Supplementary figures and images for "An extended Tudor domain within Vreteno interconnects Gtsf1L and Ago3 for piRNA biogenesis in *Bombyx mori*"

### Supplementary Figures

#### Bronkhorst et al., Figure S1

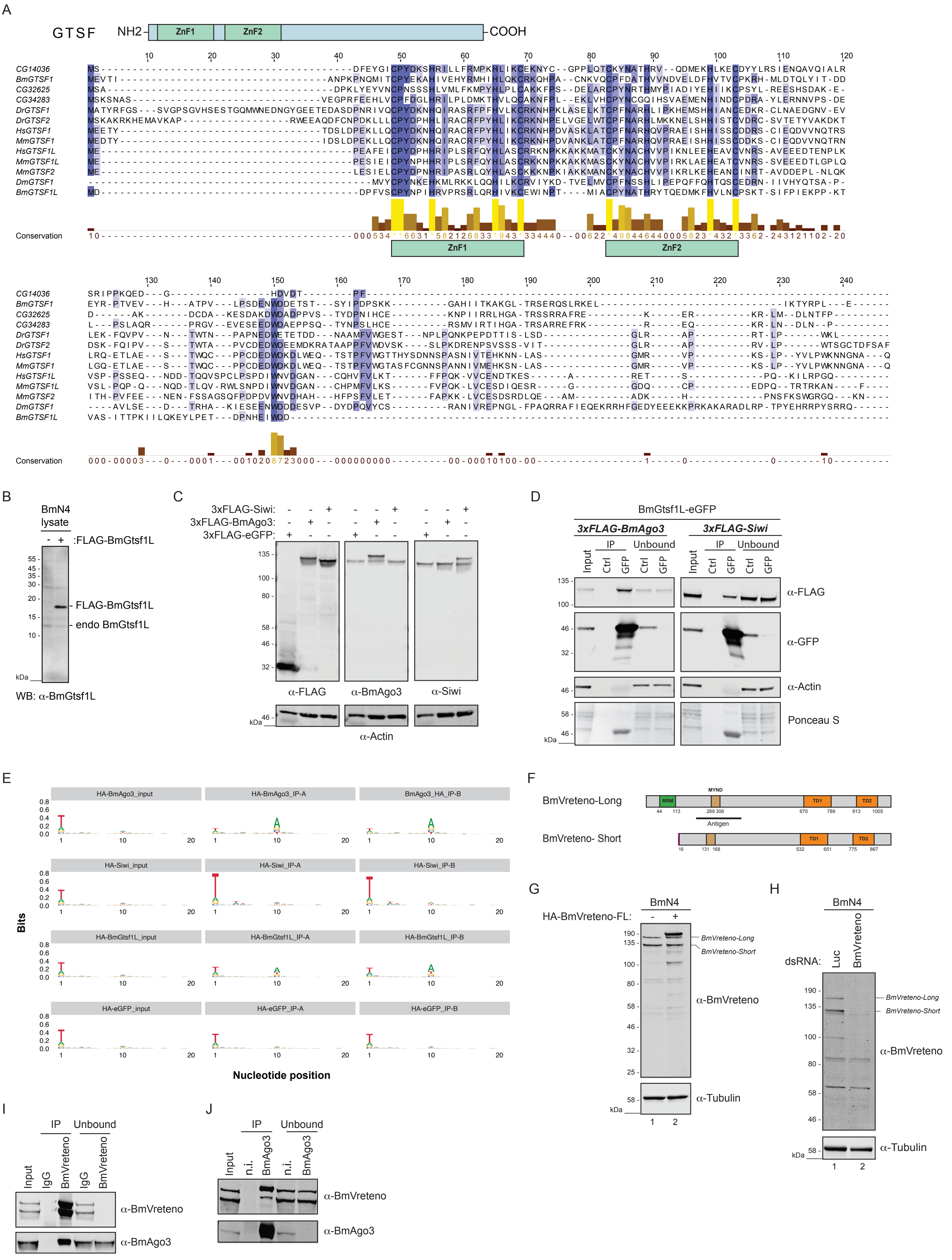

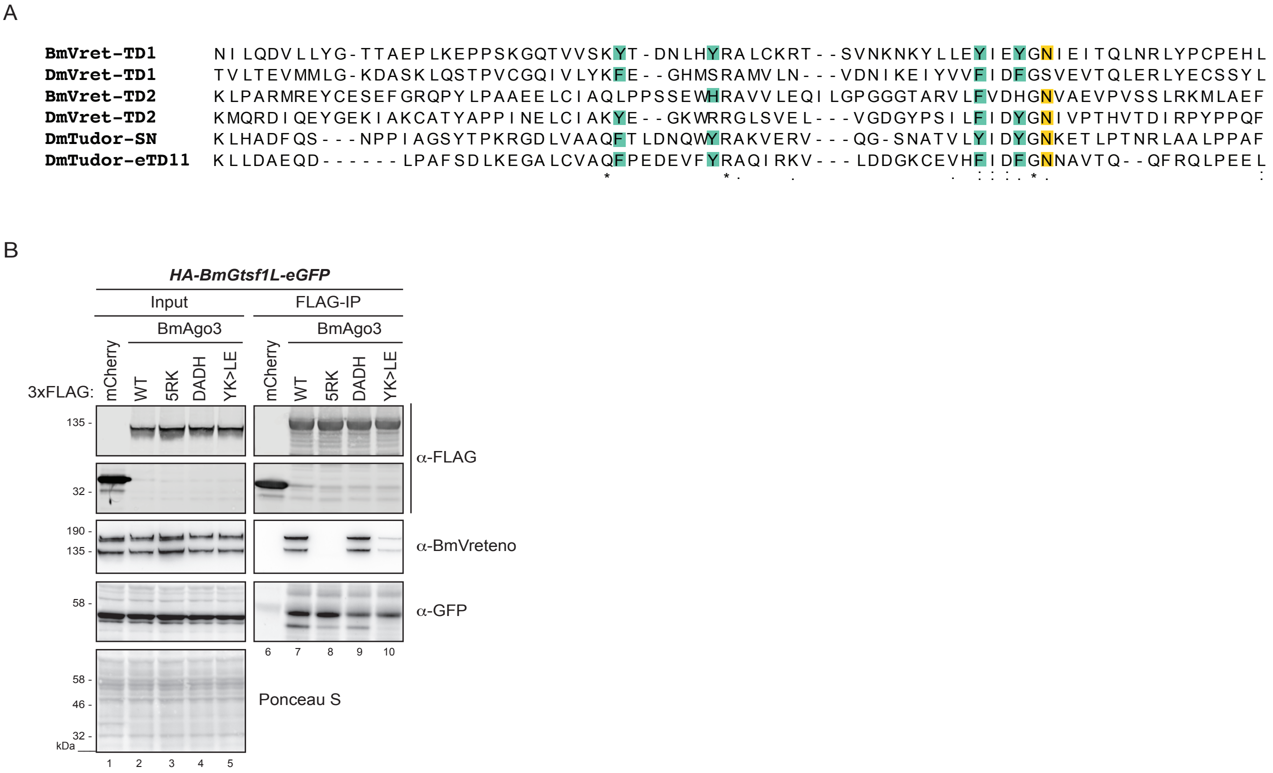

#### Bronkhorst et al., Figure S3

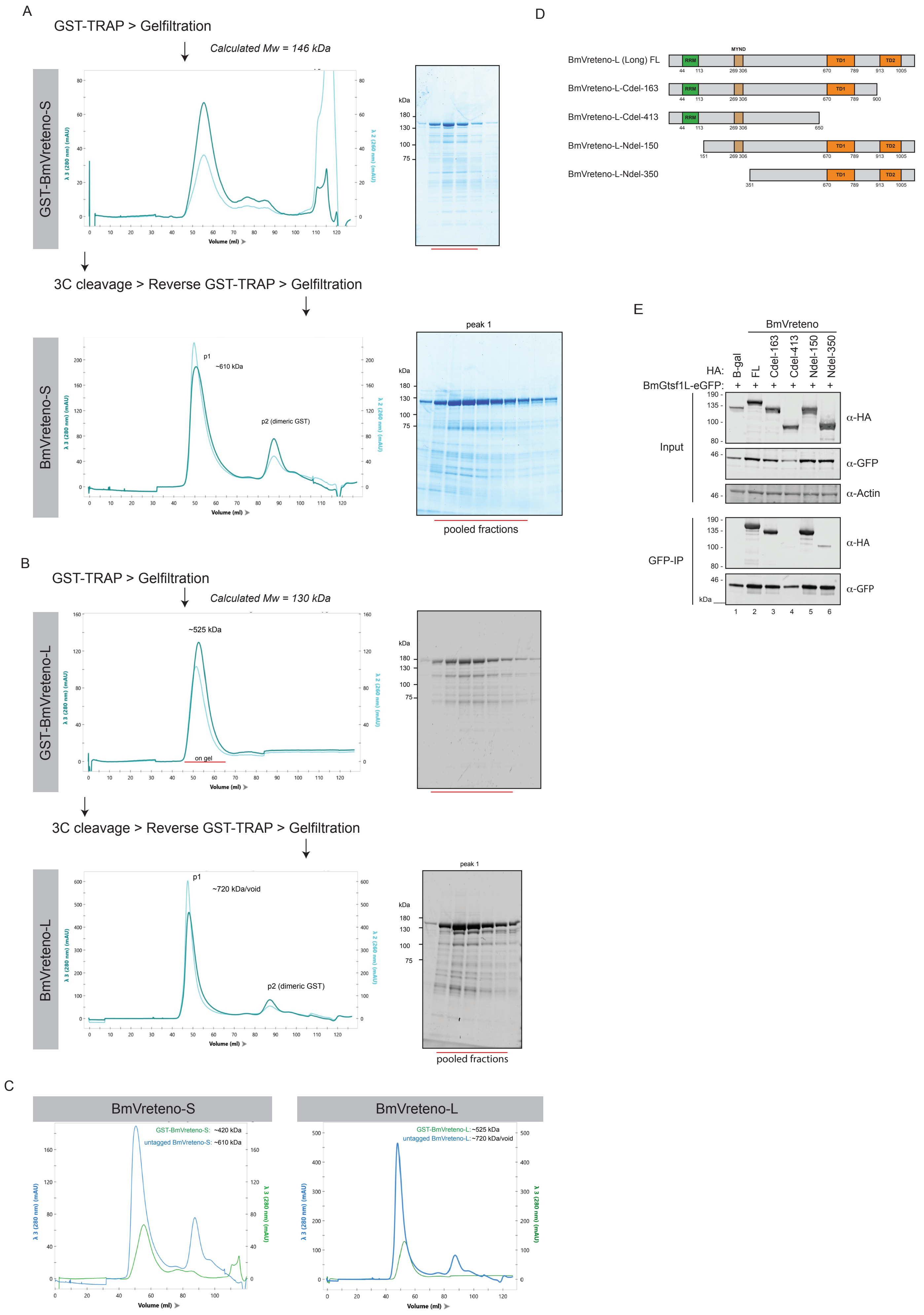

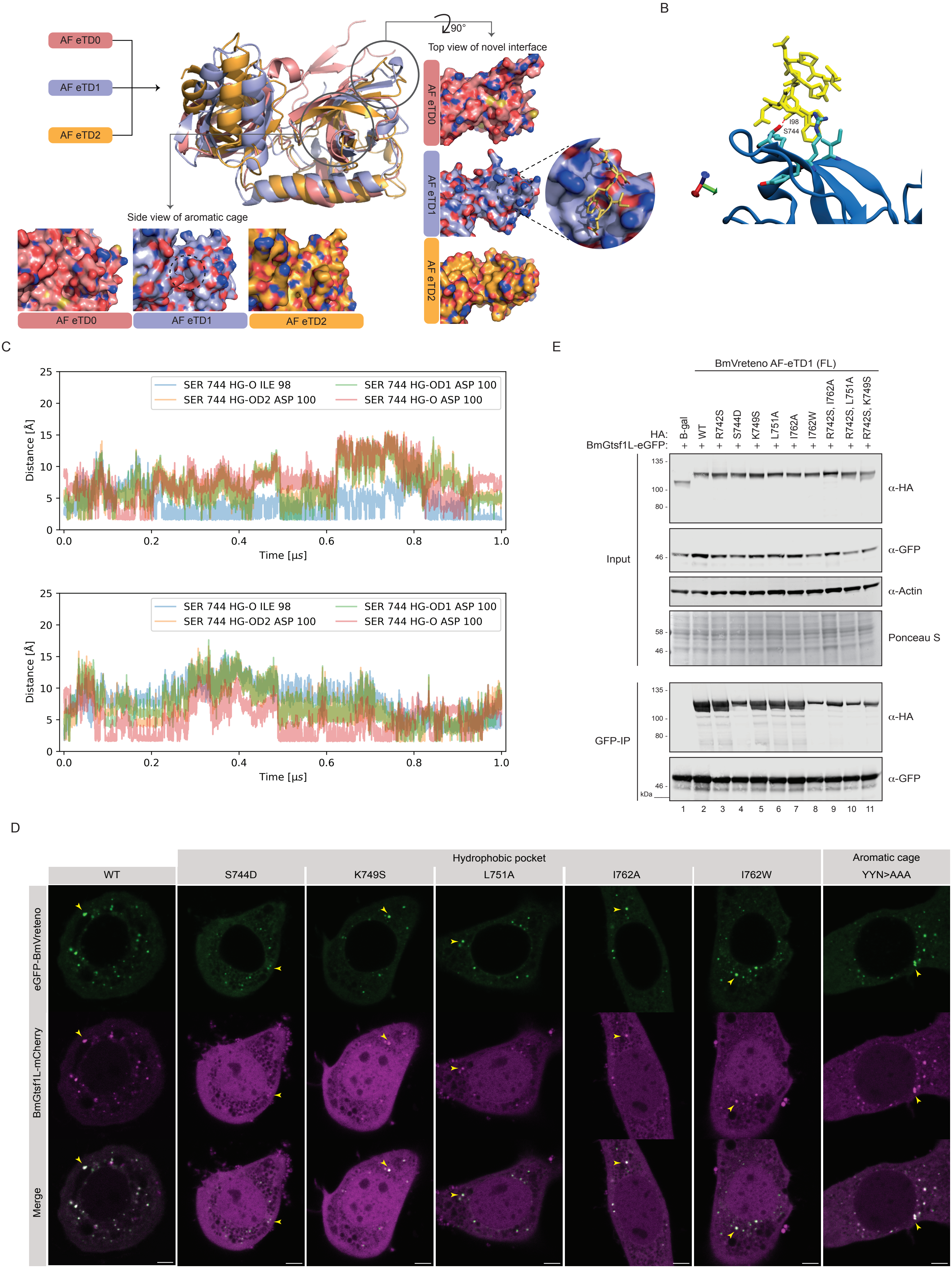
